# Prostaglandins regulate the nucleoskeleton during Drosophila border cell migration

**DOI:** 10.64898/2026.06.01.728948

**Authors:** Ashley C. Goll, Ningxin Li, Elisabeth A. Nacino, Kaden H. Bex, Sean C. Strand, Michelle S. Giedt, Tina L. Tootle

**Affiliations:** 129 E. Jefferson Street, Department of Biology, University of Iowa, Iowa City, IA 52242

**Keywords:** prostaglandins, lamin, nucleoskeleton, collective cell migration, Drosophila, border cell migration

## Abstract

The nucleoskeleton, which is comprised of Lamin A (stiffer), Lamin B, and Lamin interacting proteins, including Emerin, controls nuclear stiffness. Nuclear stiffness regulates 3D single cell migration, but its roles in collective cell migration remain unclear. To define the roles of the nucleoskeleton during collective migration we use Drosophila border cell migration. During migration the nucleoskeleton remodels. Throughout migration, Lamin A is predominantly in the nucleoskeleton of the polar cells, whereas Emerin is progressively reduced in the nucleoskeletons of both the border and polar cells, and Lamin B increases in the border cell nucleoskeleton. Further, the border cell nucleoskeleton is polarized; Lamin B is enriched in the front of the cluster while Emerin is enriched in the back. These nucleoskeletal changes require prostaglandin (PG) signaling. When PG signaling is lost, border cell migration is delayed, Lamin A and Emerin are prevalent within the border cell nucleoskeletons throughout migration and nucleoskeletal polarity is lost. Further, overexpression of Lamin A R237P in the border cells delays migration. These data reveal that border cell cluster nucleoskeletal remodeling requires PG signaling and support that this remodeling facilitates invasive, collective migration. Similar PG regulation of the nucleoskeleton likely promotes collective migration across organisms and contexts.

**Significance Statement:**

- Nucleoskeletal remodeling is critical for 3D single cell migration, but its roles in collective migration are poorly understood.
- During Drosophila border cell migration, the nucleoskeleton remodels and exhibits polarity that suggests the nuclei are softer in the front and stiffer in the back of the cluster. PG signaling is required for these nucleoskeletal changes and on-time border cell migration. Overexpression of Lamin A R237P in the border cells impairs migration.
- These results demonstrate for the first time that nucleoskeletal remodeling occurs during an *in vivo*, collective cell migration and identify PG signaling as a novel regulator of the nucleoskeleton.

## Introduction

Cell migration plays essential roles in development, wound healing, immune responses, and cancer metastasis (Merino-Casallo *et al*., 2022; Yamamoto *et al*., 2023; Cheung and Horne-Badovinac, 2025). *In vivo*, cells migrate, often as groups of cells (collective cell migration), in confined and changing three-dimensional (3D) environments (Friedl and Gilmour, 2009; Scarpa and Mayor, 2016). To understand the mechanisms regulating such migration numerous confined, 3D *in vitro* models have been developed (Liu *et al*., 2016; McGregor *et al*., 2016; Xia *et al*., 2019). Studies using these models established that the nucleus is a barrier to confined migration (Rowat *et al*., 2013; Wolf *et al*., 2013). Specifically, the nucleus must be able to deform for cells to migrate through confined spaces.

The ability of the nucleus to deform, and therefore, change its stiffness requires nucleoskeletal remodeling (Lammerding *et al*., 2005; Lammerding *et al*., 2006; Schape *et al*., 2009; Swift *et al*., 2013; Guilluy *et al*., 2014). Key nucleoskeletal components are Lamin A/C and Lamin B – which make independent type V intermediate filament networks beneath the nuclear envelope (Turgay *et al*., 2017; Tenga and Medalia, 2020) – and lamin-interacting proteins, including the Lamin A/C interactor Emerin (Mislow *et al*., 2002; Zhang *et al*., 2005; Guilluy *et al*., 2014). In general, the levels of Lamin A/C and Emerin regulate nuclear stiffness, with high levels causing stiffer nuclei (Lammerding *et al*., 2005; Lammerding *et al*., 2006; Schape *et al*., 2009; Swift *et al*., 2013; Guilluy *et al*., 2014). Thus, when Lamin B is more prevalent the nucleus is softer, facilitating the cell’s ability to maneuver through small spaces (Lammerding *et al*., 2006; Harada *et al*., 2014). Further, during confined migration, there needs to be a balance of reducing Lamin A/C and Emerin enough to allow nuclear deformation, but not to the level that results in nuclear rupture, DNA damage, and in some cases cell death (Lammerding *et al*., 2006; Davidson *et al*., 2014; Liddane *et al*., 2021). While numerous studies established that nucleoskeletal remodeling is critical for *in vitro* confined, 3D single cell migration, little is known about whether similar mechanisms are required for invasive, collective cell migration.

To address this gap in knowledge, we use border cell migration during *Drosophila* oogenesis as a model. Each *Drosophila* ovary consist of 16 to 20 ovarioles or chains of developing follicles. Each follicle is comprised of ∼650 somatic epithelial cells called follicle cells that surround the germline – 15 germline nurse cells and 1 posterior oocyte (Giedt and Tootle, 2023). During Stage 9 (S9) of oogenesis, a collective cell migration, termed border cell migration occurs (Montell, 2003; Montell *et al*., 2012; Saadin and Starz-Gaiano, 2016). At the beginning of S9, the anterior polar cells (a specialized type of follicle cell) designate 6-8 follicle cells to become border cells. These polar and border cells delaminate from the follicle cells to become the border cell cluster. This cluster migrates between the nurse cells toward the oocyte in response to secreted signaling factors from the oocyte (Duchek and Rorth, 2001; Duchek *et al*., 2001; Bianco *et al*., 2007). The polar cells remain in the center of the cluster while the border cells alter which cell is the leading cell of the cluster throughout migration (Cai *et al*., 2014). Once the border cell cluster reaches the oocyte it aids in developing the micropyle, an eggshell structure required for sperm entry and fertilization (Montell *et al*., 1992). The mechanisms regulating border cell migration are the same used to mediate collective migration across contexts, including development, wound healing and cancer (Montell, 2003; Montell *et al*., 2012; Saadin and Starz-Gaiano, 2016). Thus, this system is ideal for exploring the roles of nucleoskeletal remodeling during an *in vivo*, invasive, collective cell migration.

The key nucleoskeletal components are conserved in *Drosophila.* Indeed, flies have two lamins: the A-type lamin is Lamin C, for simplicity we refer to this as Lamin A, and the B-type lamin is Lamin Dm0, for simplicity we refer to this as Lamin B (Gruenbaum *et al*., 1988; Riemer *et al*., 1995; Schulze *et al*., 2009; Zielinska *et al*., 2025). Lamin-interacting proteins are also conserved, including Emerin (Drosophila dEmerin/Otefin; (Padan *et al*., 1990; Wagner *et al*., 2004)). One prior study examined Lamin A and Lamin B in border cell migration. This study found that Lamin B is prevalent in the border cell nucleoskeleton and contributes to delamination, maintaining nuclear integrity, and allowing the leading cell, with its protrusion, to facilitate the migration of the cluster (Penfield and Montell, 2023). As described below, our findings on the presence of lamins during border cell migration do not fully recapitulate what was previously observed. Further, whether the nucleoskeleton changes at different points in migration was not assessed in the prior study.

Here we characterize the temporal changes in the nucleoskeleton – Lamin A, Lamin B, and Emerin – throughout BC migration and identify a new regulator of nucleoskeletal remodeling during cell migration – prostaglandin (PG) signaling. PGs are short-range lipid signaling molecules that arise when arachidonic acid is acted on by cyclooxygenase (COX) enzymes to produce the PG precursor PGH_2_. PGH_2_, through the action of specific synthases, is then converted into bioactive PGs. PGs signal in an autocrine and/or paracrine fashion through G-protein coupled receptors to the activate signaling cascades and downstream targets. PGs play pivotal roles in inflammation, development, immunity, and cancer (Funk, 2001; Miller, 2006; Tootle, 2013). One function of PGs is to promote migration (Cha *et al*., 2005; Cha *et al*., 2006; Speirs *et al*., 2010; Algra and Rothwell, 2012; Rothwell *et al*., 2012). Drosophila has a single COX-like enzyme, dCOX1/Pxt (Tootle and Spradling, 2008). Prior studies show that PG signaling is required for on-time border cell migration (Fox *et al*., 2020; Mellentine *et al*., 2023). To assess the roles of PG signaling in regulating the nucleoskeleton during border cell migration we generated a new null allele of *dCOX1* and find that it recapitulates the border cell migration defects previously observed.

We find that the nucleoskeleton within the border cell cluster is dynamic during migration. Lamin A is primarily in the nucleoskeleton of the polar cells throughout border cell migration. This finding suggests the polar cell nuclei are stiffer, while the border cell nuclei are softer. The nucleoskeleton undergoes further changes during border cell migration. In early migration, both Lamin B and Emerin are prominent in the nucleoskeletons of the border and polar cells. However, as migration proceeds Lamin B increases in the border cell nucleoskeleton, whereas Emerin is reduced in the nucleoskeletons of the both the border and polar cells. These findings suggest the border cell nuclei continue to soften during migration, while the polar cell nuclei may become stiffer. We also find that the nucleoskeleton within the border cells is polarized, with Lamin B being more prominent in the nucleoskeleton of the cells at the front of the cluster and Emerin being more prominent in the back. This finding suggests that the border cells in the front of the cluster have more deformable nuclei which may facilitate the invasion of the cluster between the nurse cells; the cells at the back can retain stiffer nuclei because the nurse cells have been separated by front of the cluster. Indeed, border cells alter the topology of the nurse cells as they migrate (George *et al*., 2025).

Nucleoskeletal composition and dynamics during border cell migration are disrupted when all PG synthesis and signaling is lost. In *dCOX1* mutants, throughout migration Lamin A is present in the nucleoskeleton of both the border and polar cells. Further, when PG signaling is lost Emerin levels remains prominent in the border cell nucleoskeleton throughout migration and Lamin B levels within the border cell nucleoskeleton are reduced. The border cell nucleoskeletal polarization is also lost in the absence of PG signaling. These findings suggest the nucleoskeleton of the border cell cluster is uniformly stiffer in the absence of PG signaling, and this increased stiffness may contribute to the delays in migration observed (Fox *et al*., 2020; Mellentine *et al*., 2023). Supporting this idea, overexpression of Lamin A R237P – a mutation in Lamin A that causes rounded, and likely stiffer, nuclei (Shaw *et al*., 2022) – in the border cells causes delayed migration. Together these findings lead to the model that PG signaling drives nucleoskeletal changes within the border cell cluster that facilitate on-time migration. As both PG signaling and the nucleoskeletal components are conserved, these same mechanisms likely contribute to other collective migrations across organisms, including development, wound healing, and cancer metastasis.

## Results

### The nucleoskeletons of the border cells and polar cells are dynamic during migration

To determine whether the nucleoskeleton remains constant or is dynamic during border cell migration we stained wild-type follicles with antibodies against Lamin A, Lamin B, and Emerin and assessed different points in migration designated as early, middle, and late (Fig. 1A). We used the places within the follicle where eight nurse cells come together creating crevasses to define early, middle, and late migration (George *et al*., 2025). Early migration is from delamination at the anterior end of the follicle to immediately preceding the first nurse cell crevasse. Mid-migration is from the first crevasse to immediately preceding the second crevasse. Late migration is from the second crevasse to when the border cells reach the oocyte.

**Figure 1:**
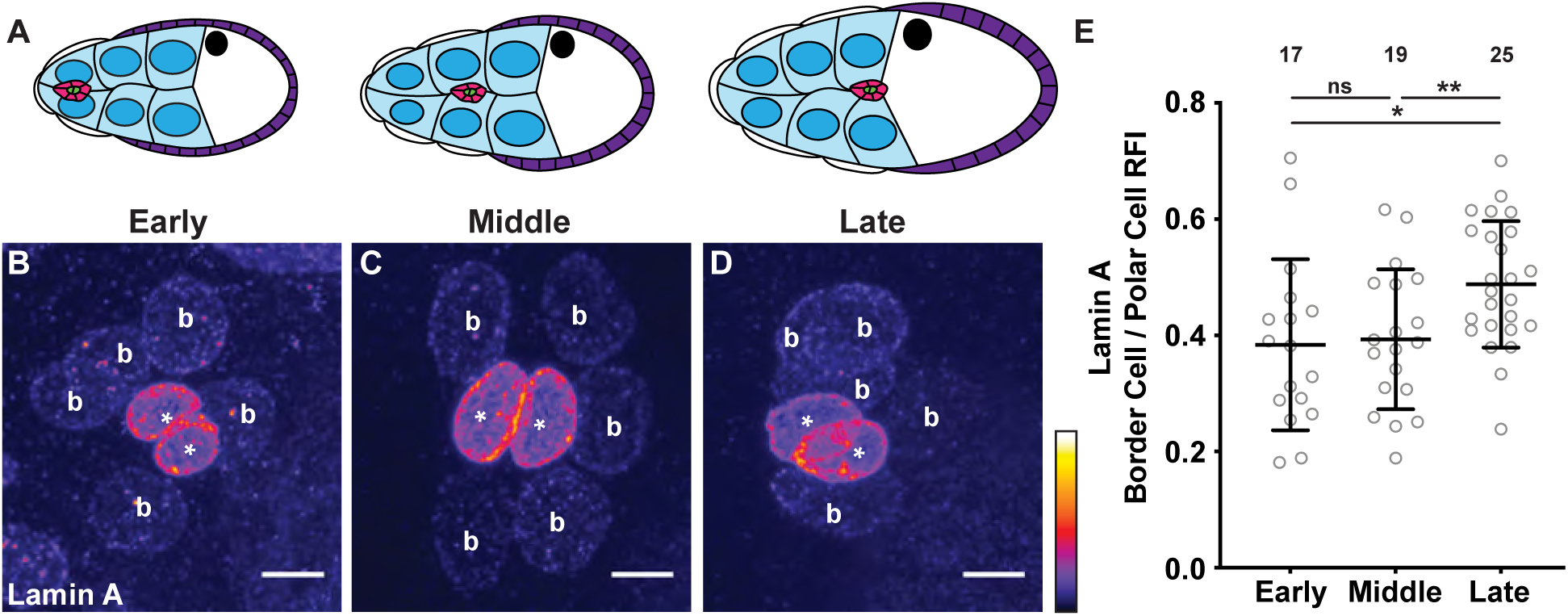
Lamin A is predominantly within the nucleoskeleton of the polar cells. **(A)** Schematic of border cell migration from early (left), middle (middle), and late (right) migration. The border cell cluster comprised of border cells (pink) and polar cells (green), nurse cells (blue), outer follicle cells (purple), stretch follicle cells (white), and oocyte (white with black nucleus) are diagramed. **(B-D)** Maximum projections of confocal slices of border cell clusters from wild-type (*yw*) S9 follicles at the indicated points in migration stained for Lamin A in Fire look up table (LUT) to reflect intensity. Border cell clusters are oriented in the direction of migration (posterior to the right). White asterisks indicate the polar cells and white b’s indicate the border cells. Images brightened by 30% to increase clarity. Scale bars = 5μm. **(E)** Graph of Lamin A border cell to polar cell relative fluorescence intensity (RFI) ratio. Circle = individual BC cluster; *n* = number of follicles, lines = averages and error bars = standard deviation (SD). ns > 0.05, ** p < 0.01, unpaired *t*-test, two-tailed. Throughout border cell migration, the nucleoskeletons of the polar cells have high, while the nucleoskeletons of the border cells have low Lamin A levels (**A-D**). This difference is reflected in the border cell to polar cell RFI of <1 at all points in migration (**E**); although there is an increase in the RFI ratio in late migration.

Lamin A appears prominent in the nucleoskeleton of the polar cells throughout migration whereas there is little Lamin A in the nucleoskeleton of the border cells (Fig. 1B-D). To quantify changes in the composition of the nucleoskeleton we measured the mean fluorescent intensity from 3D renderings of confocal image stacks using Imaris Software to obtain a relative fluorescent intensity (RFI) ratio between the border cells and polar cells (see Materials and Methods for details). There is no significant difference in the RFI ratio for Lamin A between early and mid-migration however there is an increase in the RFI ratio at late migration (Fig. 1E, p<0.05). This change in RFI could be due to increased levels in the border cell nucleoskeleton or decreased levels in the polar cells. To determine which is the case, we compared the average of the mean fluorescent intensity of the Lamin A within the nucleoskeleton of the polar cells and border cells at the different points in migration. Our data suggest the RFI increase may be due to decreased levels in the polar cells, as the level in these cells slightly decreases from mid-to late migration (SFig. 1A-B, p<0.01). We also compared the Lamin A average mean fluorescent intensity in the polar vs border cells of each individual cluster. The fluorescent intensity of Lamin A is higher in the nucleoskeleton of the polar cells in each cluster throughout migration (SFig. 1C-E; early, middle, and late p<0.0001). Together these results indicate that Lamin A is predominantly within the nucleoskeleton of the polar cells and its levels in these cells remains largely constant throughout border cell migration.

Conversely, the prevalence of both Lamin B and Emerin within the nucleoskeleton changes throughout border cell migration. Lamin B is present in the nucleoskeletons of both the polar cells and border cells during early migration. While Lamin B within the polar cells remains constant throughout migration (SFig. 2A), Lamin B prevalence in the nucleoskeleton of the border cells increases from early to late migration (Fig. 2A-C”, SFig. 2B, p<0.05). Indeed, quantification reveals a Lamin B border cell to polar cell mean RFI ratio of 0.92 during early migration and a ratio of 1.2 during late migration (Fig. 2D, p<0.001), consistent with our finding a significant increase in Lamin B within the border cell nucleoskeleton later in migration. We also compared the Lamin B average mean fluorescent intensity in the nucleoskeleton of the polar vs border cells of each individual cluster. The fluorescent intensity of Lamin B is slightly higher in nucleoskeleton of the polar cells during early and mid-migration, and in the border cells during late migration (SFig. 2C-E; early p<0.01, middle and late p<0.05). Like Lamin B, Emerin is present in the nucleoskeleton of both the polar cells and border cells during early migration (Fig. 2A’-A”). However, opposite of Lamin B, by the middle of migration Emerin appears reduced in nucleoskeleton of the border cells and by late migration there is little present (Fig. 2B’-C”). Quantification reveals that both the polar and border cells exhibit a significant decrease in Emerin prevalence from early to late migration (SFig. 3A-B, p<0.001 and p<0.0001, respectively). The Emerin border cell to polar cell mean RFI ratio is 0.72 during early migration and 0.44 during late migration (Fig. 2E, p<0.0001), consistent with there being a decrease in Emerin in the nucleoskeleton of both the border cells and polar cells as migration proceeds. We also compared the Emerin average mean fluorescent intensity in the nucleoskeleton of the polar vs border cells of each individual cluster. The fluorescent intensity of Emerin is higher in nucleoskeleton of the polar cells throughout migration (SFig. 3C-E, p<0.0001). Together, these results reveal the nucleoskeleton is remodeled during border cell cluster migration. Specifically, Emerin is reduced within the nucleoskeleton of both the border cells and polar cells, whereas Lamin B increases within the border cell nucleoskeleton as migration proceeds. These findings, along with the Lamin A findings (Fig. 1 and SFig. 1), suggest that the border cell and polar cell nuclei become softer as the cluster migrates.

**Figure 2:**
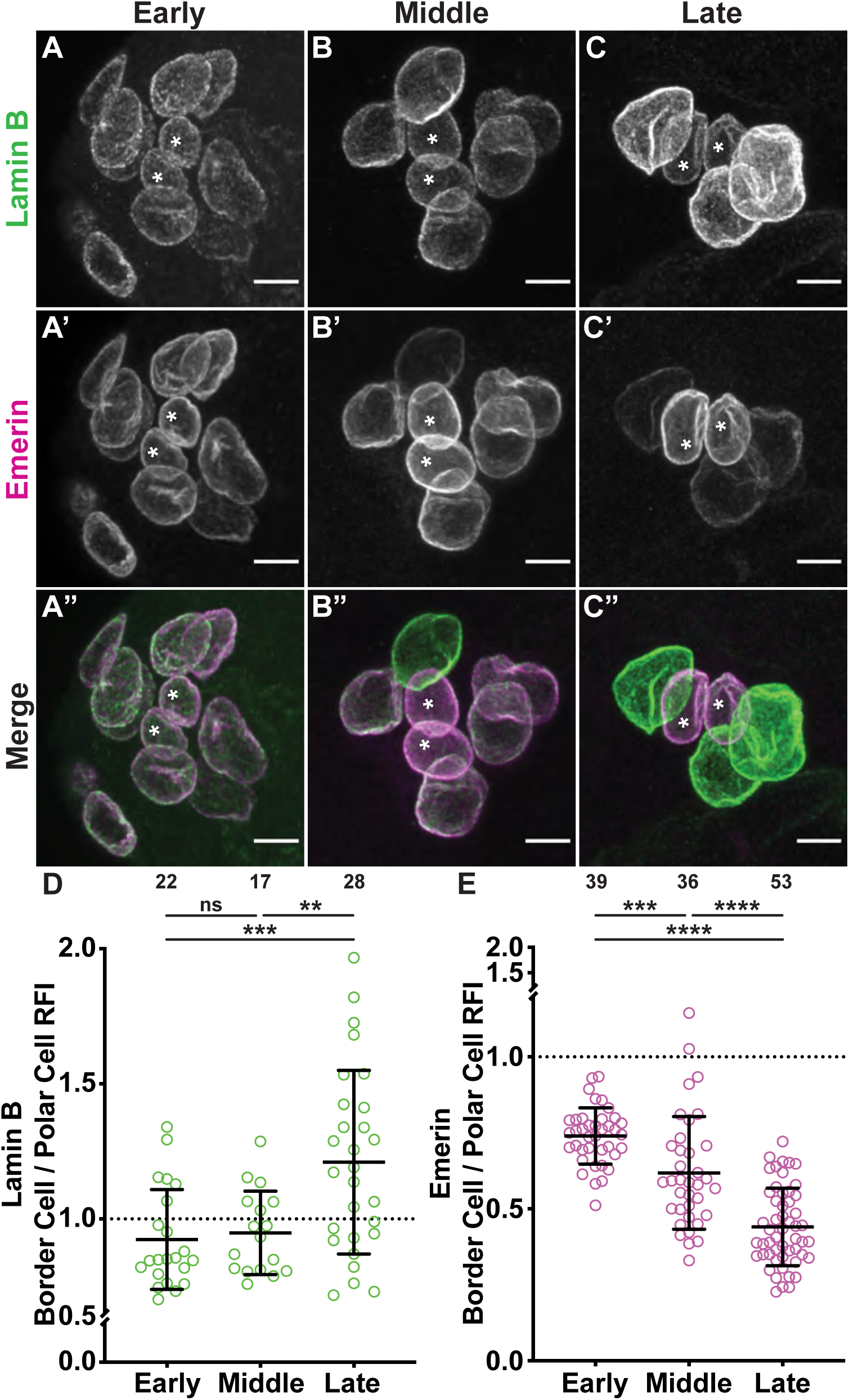
Lamin B and Emerin prevalence in the nucleoskeleton are dynamic during border cell migration. (A-C”) Maximum projections of confocal slices of border cell clusters from wild-type (*yw*) S9 follicles at the indicated points in migration stained for Lamin B (green in merge) and Emerin (magenta in merge). Border cell clusters are oriented in the direction of migration (posterior to the right). White asterisks indicate the polar cells. Images brightened by 30% to increase clarity. Scale bars = 5μm. **(D-E)**. Graphs of Lamin B (**D**) and Emerin (**E**) border cell to polar cell relative fluorescence intensity (RFI) ratio. Circle = individual BC cluster; *n* = number of follicles, lines = averages and error bars = SD. ** p < 0.01, *** p < 0.001, **** p < 0.0001 unpaired *t*-test, two-tailed. Lamin B is present in the nucleoskeletons of both the border and polar cells (**A-B”**), but by late migration Lamin B appears increased in the nucleoskeleton of the border cells compared to the polar cells (**B-C”**); this increase is reflected in the RFI >1 during late migration (**D**). Emerin is also initially present in the nucleoskeletons of both the border and polar cells (**A’-A”**) but is progressively reduced in the nucleoskeleton of the border cells as migration proceeds (**B’-C”**); this change is reflected in the reduced RFI in late migration (**E**).

### The border cell nucleoskeleton is polarized during migration

We also find that Lamin B and Emerin exhibit a polarized accumulation within the nucleoskeleton of the border cells during migration. Specifically, Lamin B appears more prevalent in the nucleoskeleton of the border cells at the front of the cluster compared to those at the back, while Emerin exhibits the opposite pattern and appears to be at higher levels in the nucleoskeleton at the back of the cluster (Fig. 3A-B”). To quantify this polarized localization, we subtracted the average mean fluorescent intensity of the back border cells from the front border cells of each cluster. A positive value indicates that the nucleoskeletal protein is more prominent at the front of the border cell cluster whereas a negative value indicates that the nucleoskeletal protein is more prominent at the back of the cluster. The mean fluorescent intensity difference for Lamin B is 2.2 while the difference for Emerin is -1.8 (Fig. 3C-D), supporting the nucleoskeleton is polarized. Lamin B prominence at the front of the cluster is consistent throughout early, middle, and late migration as all mean fluorescent intensity differences are positive (SFig. 4A; 3.01, 1.8, and 2.0, respectively). Similarly, Emerin prominence at the back of the cluster is consistent during early and mid-migration but is more uniform during late migration due to the decreased levels in the border cells described above (SFig. 4B; -2.4, -2.1, and -0.79, respectively). Taken together these results suggest that the nuclei at the front of the cluster are softer than those at the back of the cluster.

**Figure 3:**
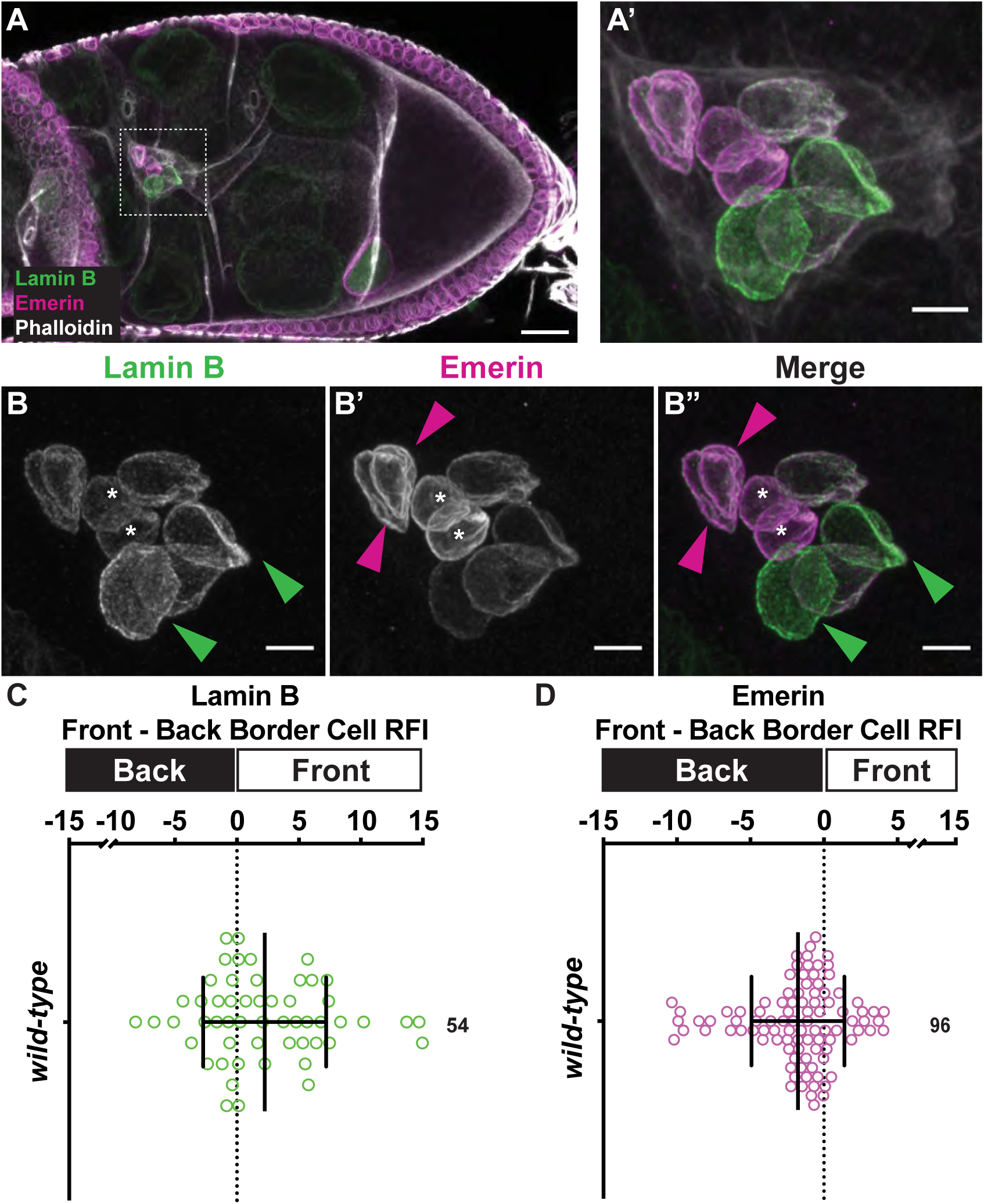
Lamin B and Emerin are polarized within the border cell cluster. **(A)** Maximum projection of confocal slices of a wild-type (*yw*) S9 follicle stained for Lamin B (green in merge), Emerin (magenta in merge), and F-actin (phalloidin, white in merge). Scale Bar = 20 μm. **(A’-B”)** Zoomed in maximum projections of confocal slices of the white boxed region in **A**. Border cell clusters are oriented in the direction of migration (posterior to the right). White asterisks indicate the polar cells. In **B-B”**, green arrowheads indicate border cells where Lamin B is more prominent and magenta arrowheads indicate border cells where Emerin is more prominent. Images brightened by 30% to increase clarity. Scale bars = 5μm. **(C-D)** Graphs of front - back relative fluorescent intensity (RFI) ratio for Lamin B (**C**) and Emerin (**D**). Circle = individual BC cluster; *n* = number of follicles, lines = averages and error bars = SD. A value of 0 indicates a lack of polarity, represented by the dotted line. Positive value indicates the nucleoskeletal protein is more prominent at the front of the cluster. Negative value indicates nucleoskeletal protein is more prominent at the back of the cluster. Lamin B is prominent at the front of the cluster (**A-B”, C**). Emerin is prominent at the back of the cluster (**A-B”, D**).

### PGs are required for nucleoskeletal remodeling during border cell migration

Having found the nucleoskeleton is dynamic and exhibits polarity during border cell migration, we next sought to identify mechanisms regulating the nucleoskeleton during border cell migration. As prior data uncovered PGs are required for on-time border cell migration (Fox *et al*., 2020; Mellentine *et al*., 2023), we hypothesized that PGs may regulate nucleoskeletal dynamics to facilitate border cell cluster migration. To test our hypothesis, CRISPR was used to fully excise the *dCOX1* gene (see Materials and Methods), resulting in a null allele (SFig. 5); we refer to this as *dCOX1-/-*. Loss of dCOX1 results in the complete loss of PG synthesis and signaling (Tootle and Spradling, 2008). We verified that like the previously assessed transposon insertional alleles of *dCOX1*, the new null allele results in border cell defects during S9. Specifically, we assessed border cell migration by confocal microscopy and our migration index quantification (Fox *et al*., 2020; Lamb *et al*., 2020; Mellentine *et al*., 2023). To quantify the migration index, the distance the border cell cluster has traveled from the anterior end of the follicle is divided by the distance the outer follicle cells have traveled from the anterior end of the follicle. If the ratio is ∼1 it indicates that the border cell cluster is migrating on time, as the border cells normally remain in-line with the position of the outer follicle cells throughout migration. If the ratio is <1 it indicates a delay in migration. As expected, in wild-type follicles the border cell cluster is largely in-line with the outer follicle, while migration is delayed in *dCOX1-/-* follicles (SFig. 6A, migration index of 0.74 compared to 0.89, p<0.0001). We also measured the length of the border cell cluster and, as previously observed, the *dCOX1-/-* clusters are significantly longer than wild-type clusters (SFig. 6B, 28µm compared to 24µm, p<0.0001). There is also an increase in the number of border cells when PG signaling is lost. In wild-type follicles there is an average of ∼5 border cells while in *dCOX1-/-* follicles there is an average of ∼7 border cells per cluster (SFig. 6C, p<0.001). Thus, the new *dCOX1* allele recapitulates the border cell phenotypes observed with the previously studied alleles (Fox *et al*., 2020).

We used this *dCOX1* allele to determine whether PG signaling regulates the nucleoskeleton during border cell migration. We first assessed Lamin A. Unlike wild-type follicles in which Lamin A is primarily restricted and at high levels within the polar cell nucleoskeleton (Fig. 4A-C), when PG signaling is lost, Lamin A remains high within the border cell nucleoskeleton (Fig. 4D-F). Quantification of the Lamin A border cell to polar cell mean RFI ratio for wild type is 0.40 while *dCOX1-/-* has a RFI ratio of 0.48, consistent with the stronger prevalence of Lamin A in the border cells when PG signaling is lost (Fig. 4G, p<0.01). To gain more insight into the changes in Lamin A within the cluster when PGs are lost, we quantified the mean fluorescent intensity of the polar cells and border cells separately. Loss of dCOX1 results in a higher prevalence of Lamin A within the nucleoskeletons of both the polar and border cells (SFig. 7A-B, p<0.0001). These data indicate that PG signaling is required to limit Lamin A presence in the nucleoskeletons of the polar cells and border cells during migration.

**Figure 4:**
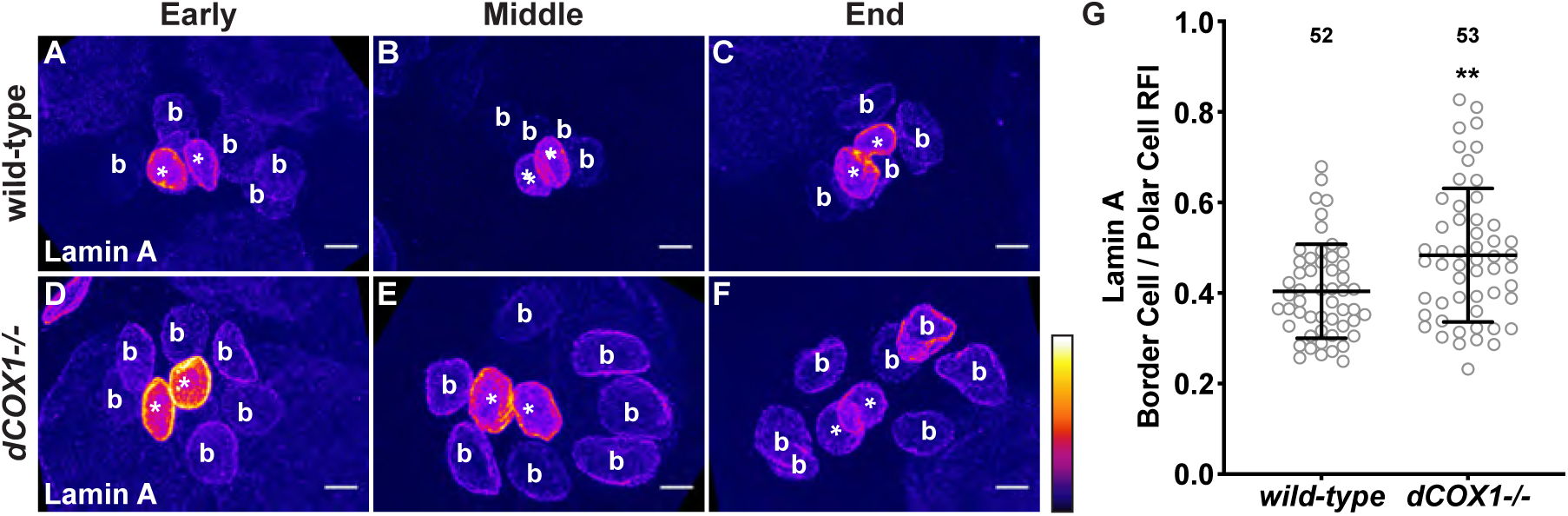
PG signaling restricts Lamin A predominantly to the nucleoskeleton of the polar cells. (A-F) Maximum projections of confocal slices of border cell clusters from wild-type (*yw*, **A-C**) and *dCOX1-/-* (**D-F**) S9 follicles at the indicated points in migration stained for Lamin A in Fire LUT to reflect intensity. Border cell clusters are oriented in the direction of migration (posterior to the right). White asterisks indicate the polar cells and white letter b’s indicated the border cells. Images brightened by 30% to increase clarity. Scale bars = 5μm. **(G)** Graph of Lamin A border cell to polar cell relative fluorescence intensity (RFI) ratio. Circle = individual BC cluster; *n* = number of follicles, lines = averages and error bars = SD. ** p < 0.01, unpaired *t*-test, two-tailed. Lamin A is at a higher level in the nucleoskeleton of the border cells of *dCOX1-/-* (**D-F**) compared to wild-type follicles (**A-C**), resulting in a higher RFI (**G**).

We next assessed if PG signaling regulates Lamin B and Emerin during border cell migration. Lamin B appears largely unaffected by the loss of PG signaling. Specifically, Lamin B is present in the polar cells and border cells throughout migration in both wild-type and *dCOX1-/-* (Fig. 5A-F). Quantification of the mean fluorescent intensity reveals that the Lamin B prevalence in the nucleoskeleton of the polar cells is similar in both wild-type and *dCOX1-/-* (SFig. 8A). However, when Lamin B border cell fluorescent mean intensity is compared, the prevalence of Lamin B in wild-type follicles is trending slightly higher than in *dCOX1-/-* (SFig. 8B, p=0.061). Indeed, quantification of the Lamin B border cell to polar cell mean RFI ratio indicates a significant increase in Lamin B in the nucleoskeleton of wild-type compared to *dCOX1-/-* follicles (Fig. 5G p<0.01). These results suggest that PG signaling normally promotes Lamin B accumulation within the nucleoskeleton of the border cells.

**Figure 5:**
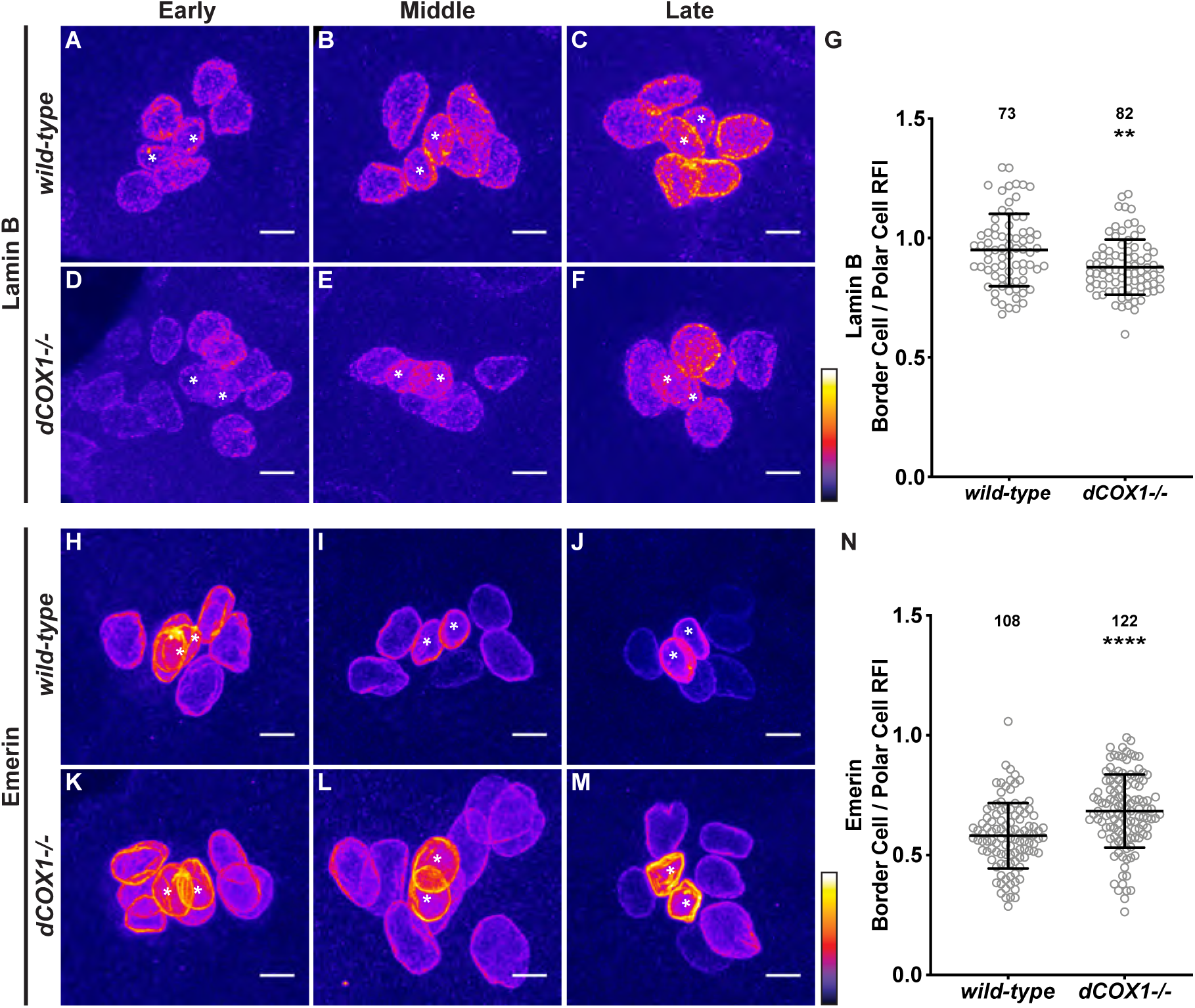
PG signaling promotes the increase in Lamin B and the removal of Emerin from the nucleoskeleton of the border cells during migration. (A-F, H-M) Maximum projections of confocal slices of border cell clusters from wild-type (*yw*; **A-C, H-J**) and *dCOX1-/-* (**D-F, K-M**) S9 follicles at the indicated points in migration stained for Lamin B (**A-F**) or Emerin (**H-M**) in Fire LUT to reflect intensity. Border cell clusters are oriented in the direction of migration (posterior to the right). White asterisks indicate the polar cells. Images brightened by 30% to increase clarity. Scale bars = 5μm. **(G, N)** Graph of border cell to polar cell relative fluorescence intensity (RFI) ratio for Lamin B (**G**) and Emerin (**N**). Circle = individual BC cluster; *n* = number of follicles, lines = averages and error bars = SD. ***p<0.001, **** p < 0.0001, unpaired *t*-test, two-tailed. **(H-I)**. Lamin B appears to be more prevalent in the nucleoskeleton of the border cells of wild-type follicles (**A-C**) compared to *dCOX1-/-* (**D-F**); this is reflected in the lower RFI ratio when PG signaling is lost. Emerin remains at a high level within the nucleoskeleton of the border cells in *dCOX1-/-* follicles (**K-M**) compared to wild-type (**H-J**), resulting in a higher RFI ratio when PG signaling is lost (**N**).

Similar to what we observed for Lamin A, loss of PG signaling results in Emerin remaining present in the nucleoskeleton of the border cells throughout migration (Fig. 5K-M); in wild-type follicles Emerin is at higher levels in the nucleoskeleton of the polar cells than the border cells, but decreases in both cell types during migration (Fig. 5H-J). Quantification of the Emerin border cell to polar cell mean RFI ratio reveals that loss of PGs results in a RFI ratio of 0.68 compared to 0.58 in wild-type (Fig. 5N, p<0.0001), consistent with the appearance of a higher prevalence of Emerin in the nucleoskeleton of the border cells in *dCOX1-/-*. To further assess the changes in Emerin within the cluster when PGs are lost, we quantified the mean fluorescent intensity of the polar cells and border cells separately. Loss of dCOX1 results in a higher prevalence of Emerin within the nucleoskeletons of both the polar and border cells (SFig. 8C-D; polar cell p<0.01 and border cell p<0.0001). Together these results indicate that PG signaling limits the prevalence of Emerin in the nucleoskeleton of the border cell cluster.

In summary, loss of PGs results in an increase in Lamin A and Emerin in the nucleoskeleton of all cells within the border cell cluster and a mild decrease in Lamin B in the border cells. These findings suggest that PG signaling is required for the nucleoskeleton of the border cell cluster to remodel and likely soften.

### PGs regulate the nucleoskeletal polarity of the border cell cluster

PG signaling also regulates nucleoskeletal polarity within the border cell cluster during migration. Unlike wild-type follicles, in which the border cells have a higher nucleoskeletal prevalence of Lamin B at the front of the cluster and Emerin at the back of the cluster, in *dCOX1-/-* follicles this polarity is lost. Lamin B and Emerin appear more uniform throughout the border cell cluster in the absence of PG signaling (Fig. 6A-B”). As in Figure 3, to quantify the polarity we subtracted the mean fluorescent intensity of the border cells at the back of cluster from the mean fluorescent intensity of the cells at the front. The mean fluorescent intensity difference for Lamin B in wild-type follicles is 2.7, whereas for *dCOX1-/-* follicles it is 0.28 (Fig. 6C, p<0.01), and for Emerin it is -1.94 in wild-type and -0.24 in *dCOX1-/-* (Fig. 6D, p<0.01). These results indicate that Lamin B and Emerin are largely evenly present in the nucleoskeleton of the border cells throughout the cluster when PG signaling is lost. Together our findings suggest that in the absence of PG signaling the nuclei of the border cell cluster are stiffer. The increased stiffness of the border cells, particularly at the front of the cluster, may impede the ability of the ability of the nuclei to deform during migration. Thus, the nucleoskeletal changes when PG signaling is lost could contribute to the delayed migration observed (SFig. 6A).

**Figure 6:**
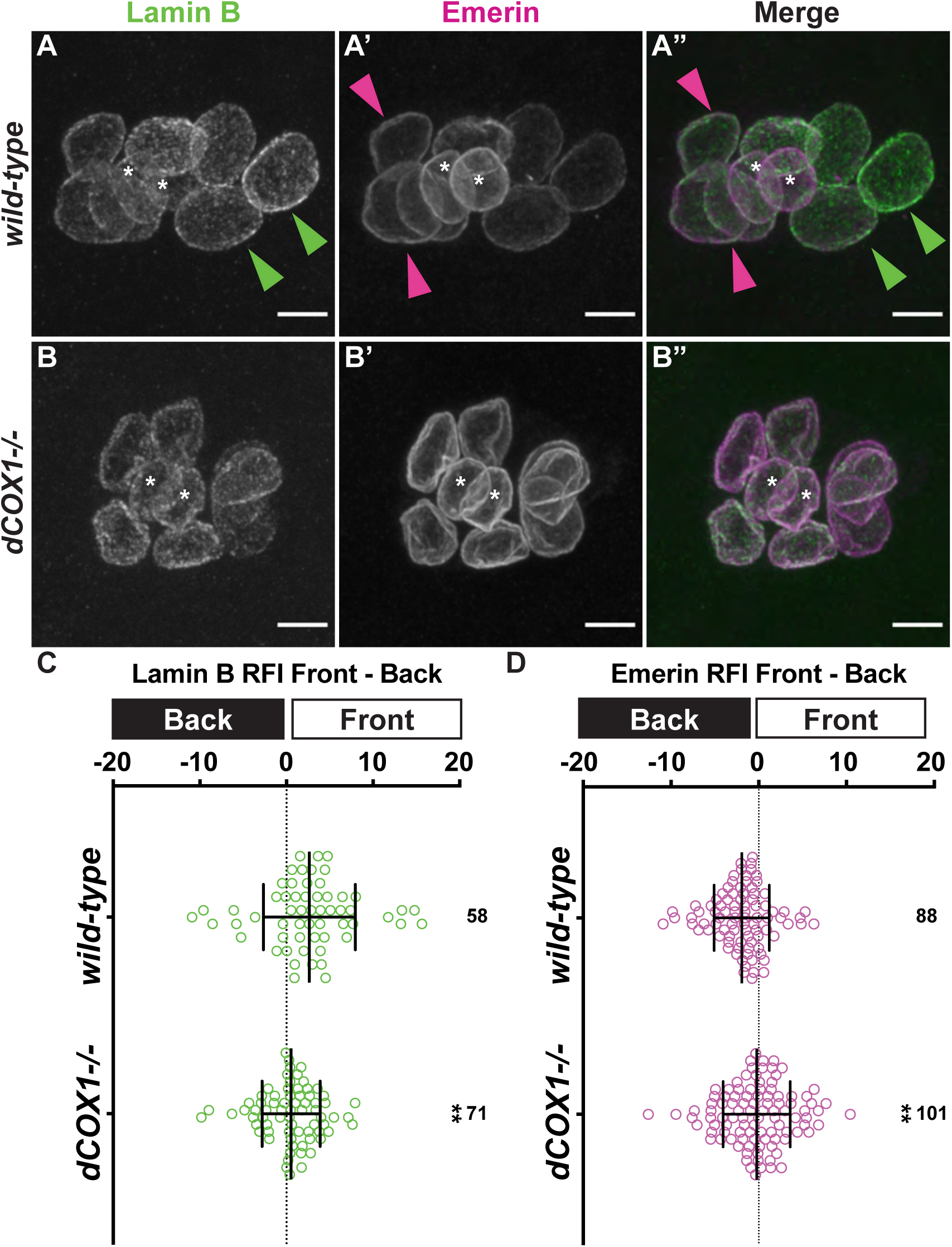
PG signaling is required for nucleoskeletal polarity within the border cell cluster. (A-B”) Maximum projections of confocal slices of border cell clusters from wild-type (*yw*, **A-A”**) and *dCOX1-/-* (**B-B”**) S9 follicles stained for Lamin B (green in merge) and Emerin (magenta in merge). Border cell clusters are oriented in the direction of migration (posterior to the right). White asterisks indicate the polar cells. Green arrowheads indicate border cells where Lamin B is more prominent. Magenta arrowheads indicate border cells where Emerin is more prominent. Images brightened by 30% to increase clarity. Scale bars = 5μm. **(C-D)** Graph of front - back relative fluorescent intensity (RFI) ratio for Lamin B (**C**) and Emerin (**D**). Circle = individual BC cluster *n* = number of follicles, lines = averages and error bars = SD, ** p <0.01, unpaired *t*-test, two-tailed. A value of 0 indicates a lack of polarity, represented by the dotted line would. Positive value indicates the nucleoskeletal protein is more prominent at the front of the cluster. Negative value indicates nucleoskeletal protein is more prominent at the back of the cluster. Loss of PG signaling results in Lamin B and Emerin levels within the nucleoskeleton of the border cells being similar in the front and back of the cluster (**B-D**), whereas Lamin B is more prominent in the front and Emerin in the back of the cluster in wild-type follicles (**A-A”, C-D**).

### Increasing Lamin A R237P levels within the border cell nucleoskeleton delays migration

Our finding that Lamin A is prominent within the nucleoskeleton of the border cells when PG signaling is lost suggests this change in nucleoskeletal composition, and likely nuclear stiffness, may contribute to migration delays. If this is the case, then overexpression of Lamin A in the border cells should similarly impair migration. We used the UAS/GAL4 system to overexpression Lamin A in the border cells and find, unexpectedly, that it has no impact on border cell migration (SFig. 9, overexpression migration index (MI) = 1.009; GAL4 Only MI = 1.003; UAS only MI = 0.97). This lack of a phenotype could mean that our hypothesis is wrong or that Lamin A levels were not sufficiently increased. To further test the role of Lamin A in border cell migration, we overexpressed a Lamin A mutant in the border cells, R237P. This mutation is a fly model of a human mutation (H222P) observed in Emery Dreifuss Muscular Dystrophy (Bonne *et al*., 2000). Prior work with R237P in larval muscles revealed it localizes to the nucleoskeleton; it exhibits rounded nuclei, suggestive of increased nuclear stiffness; and the nuclei remain mechanically responsive (Shaw *et al*., 2022). These findings led us to speculate that overexpression of R237P in the border cells will increase Lamin A levels in the nucleoskeleton and drive nuclear stiffening. Indeed, we find overexpression of R237P in the border cells, increases Lamin A in the border cells (Fig. 7A-C’), and causes delayed migration (Fig. 7D, MI of 0.85 compared to 1.00 and 1.06, p<0.0001). This finding supports that model that the nucleoskeletal composition of the border cells is critical for on-time migration; too much Lamin A, and thus likely stiffer nuclei, impedes the invasive, collective migration of the border cells.

**Figure 7:**
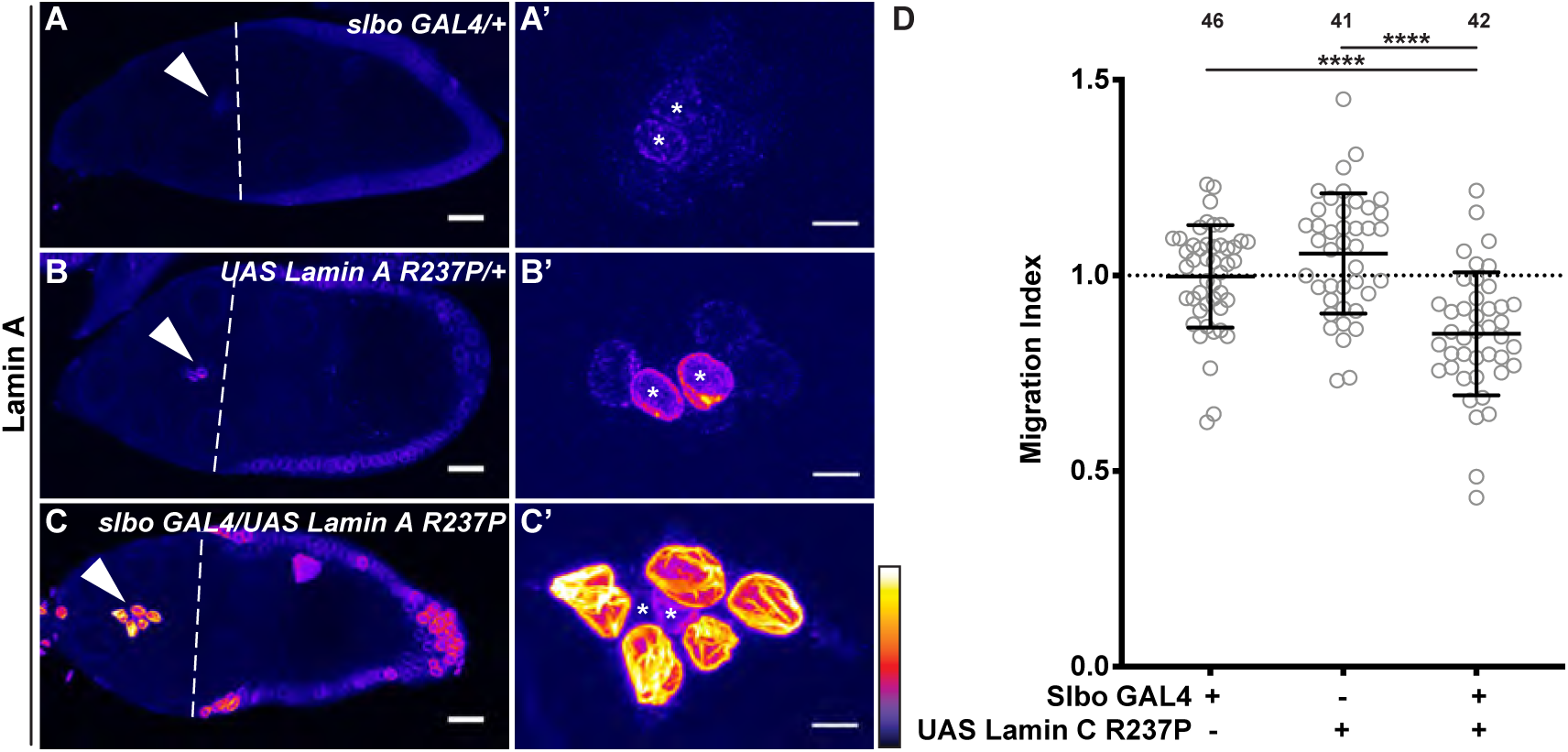
Overexpression of Lamin A R237P delays border cell migration. (A-C) Maximum projection of confocal slices of S9 follicle of the indicated genotypes stained for Lamin A in Fire LUT to reflect intensity. White arrow heads indicate border cell cluster and white dashed lines indicate the outer follicle cell position. Scale bars= 20 μm. **(A’-C’)** Zoomed in maximum projection of the border cell clusters from the follicles in A-C. Border cell clusters are oriented in the direction of migration (posterior to the right). White asterisks indicate the polar cells. Scale bars = 5μm. Images brightened by 30% to increase clarity. (**A-A’**) *slbo GAL4/+*; (**B-B’**) *UASt Lamin A R237P/+;* (**C-C’**) *UASt Lamin A R237P/+; slbo GAL4/+*. **(D)** Graph of migration index for the indicated genotypes. On-time border cell migrated indicated by dotted line. Circle = individual BC cluster; *n* = number of follicles, lines = averages and error bars = SD. **** p < 0.0001, unpaired *t*-test, two-tailed. Both controls exhibit on-time migration (**A-B’, D**), while overexpression of Lamin A R237P in the border cells delays migration (**C-D**).

## Discussion

Using Drosophila border cell migration as an *in vivo* model of invasive, collective cell migration we provide some of the first evidence that nucleoskeletal changes facilitate collective migration and identify PG signaling as a novel regulator of the nucleoskeleton. While Lamin A is primarily restricted to the nucleoskeleton of the polar cells throughout border cell migration, Lamin B and Emerin levels and cell-type prevalence change during migration. Specifically, early in migration, Lamin B and Emerin are present within the nucleoskeletons of both the border and polar cells. As migration proceeds, Lamin B levels increase in the nucleoskeleton of the border cells. Whereas Emerin levels decrease in the nucleoskeletons of both the border and polar cells; Emerin, at all stages of migration, is higher in the polar cells. Together, these findings suggest that both the border and polar cell nuclei soften throughout migration. However, the polar cell nuclei are stiffer than the border cells. The nucleoskeleton of the border cells also exhibits polarity within the cluster. During early and mid-migration, when both Lamin B and Emerin are prevalent in the nucleoskeleton of the border cells, they exhibit front/back polarity; Lamin B is high in the border cells at the front, while Emerin is high at the back of the cluster. These findings suggest the cells at the front of the cluster exhibit softer nuclei, while those at the back are stiffer.

PG signaling promotes both nucleoskeletal remodeling and polarity during border cell migration. Loss of dCOX1 results in Lamin A and Emerin being prevalent in the nucleoskeleton of both the border and polar cells throughout migration, whereas Lamin B is reduced in the nucleoskeleton of the border cells. Further, nucleoskeletal polarity is lost in the absence of PG signaling. Together, the finding that nucleoskeletal changes during border cell migration require PG signaling, along with the finding that PG signaling is required for on-time border cell migration (Fox *et al*., 2020; Mellentine *et al*., 2023), suggest that the nucleoskeletal changes are needed to facilitate on-time migration. Supporting this model, overexpression of a Lamin A mutant, R237P (Shaw *et al*., 2022), in the border cells delays migration. Together these data lead to the model that PG signaling drives nucleoskeletal remodeling necessary for on-time border cell migration. As PGs are widely implicated in cell migration (Cha *et al*., 2005; Cha *et al*., 2006; Li *et al*., 2012; Menter and Dubois, 2012), this same mechanism is likely used in other migratory contexts, including cancer metastasis.

### Nucleoskeleton and confined migration

Studies over the past two decades established that nuclear deformation during confinement and the ability of cells to undergo 3D migration depend on the nucleoskeletal composition and thereby, the ability of the nucleus to deform (Wolf *et al*., 2013; Liu *et al*., 2016; McGregor *et al*., 2016; Xia *et al*., 2019). The reduction in Lamin A/C and Emerin must be tightly regulated to facilitate nuclear deformation, but not so reduced that it results in nuclear rupture, DNA damage, and/or cell death. For example, in the context of confinement – Lamin A/C but not Lamin B – regulates nuclear stiffness. Loss of Lamin A/C results in reduced stiffness and increased nuclear fragility, leading to increased cell death under mechanical strain (Lammerding *et al*., 2006). Microfluidic experiments uncovered the level of Lamin A/C determines the cells ability to migrate through narrow constrictions, as reduced Lamin A/C allowed faster migration (Davidson *et al*., 2014). Similarly, Emerin levels must be tightly regulated during migration in micropores and invasion, as overexpression of Emerin in invasive breast cancer cells impairs migration (Liddane *et al*., 2021). While these and other studies established that *in vitro*, nucleoskeletal changes control nuclear stiffness and the ability of cells to migrate through narrow 3D environments, the importance of this to *in vivo*, non-disease cell migrations remained unclear. Among the first studies to support the physiological relevance of controlling nuclear stiffness during cell migration, Rowat *et al*. showed that neutrophils, which have small, irregularly shaped nuclei and migrate through some of the tightest spaces *in vivo*, tightly limit Lamin A levels to migrate through micro-scale pores (Rowat *et al*., 2013). Together these studies reveal that nucleoskeletal composition is critical for single cells to migrate in confined 3D environments.

While nucleoskeletal changes are necessary for confined migration *in vitro*, few studies have looked at the role of the nucleoskeleton *in vivo* or during development. Here we show that the nucleoskeletal changes during Drosophila border cell migration mirror those seen for confined, single cell migration; in the migrating border cells, Lamin A and Emerin are reduced but Lamin B is retained within the nucleoskeleton. This work is among the first assessments of nucleoskeletal dynamics during an *in vivo*, invasive collective migration. However, evidence supports that similar nucleoskeletal regulation occurs during other developmental and physiological migrations. For example, in *C. elegans*, hypodermal P-cell migration requires nuclear deformation that depends on a functional Linker of the Nucleoskeleton and Cytoskeleton (LINC) Complex, suggesting that force transmission to the nucleus is essential for this confined migration (Bone *et al*., 2016). However, due to tool limitations they were unable to assess changes in the nuclear lamina. In Drosophila embryos, macrophages invasively migrate into the germband. While RNAi knockdown of either Lamin A or B did not have an effect on their own, when migration is impaired, reducing either lamin rescues invasion; this suggests the nucleus can impede macrophage migration (Belyaeva *et al*., 2022). Drosophila hemocytes, which are analogous to leukocytes, migrate within narrow vessel-like structures in the pupal wing and can invade the tissue in response to damage (Karling and Weavers, 2025). These hemocytes exhibit high levels of Lamin B, but little to no Lamin A, similar to what we find in the border cells. Upon migrating into a constriction, Lamin B levels increase. Reducing either lamin impaired nuclear deformation and migration, and resulted in nuclear rupture (Karling and Weavers, 2025). In a recent preprint, zebrafish neural crest cells exhibit nuclear deformations that scale with tissue confinement; this work reveals that during confinement the cells decrease Lamin B2 levels, allowing the nuclei to stretch and not rupture, and thereby, impeding DNA damage (Häkkinen *et al*., 2025). Thus, there is growing evidence that nucleoskeletal changes driving nuclear softening are required for *in vivo*, confined, cell migrations.

### The nucleoskeleton is remodeled during border cell migration

We find the nucleoskeleton undergoes dynamic changes throughout border cell migration. Lamin A is predominantly within the nucleoskeleton of the polar cells at all points in migration but exhibits a decrease in its level in late migration. This finding is not consistent with a previous study on lamins during border cell migration. Specifically, Penfield and Montell report Lamin A is present in the nucleoskeleton of the border cells during migration (Penfield and Montell, 2023). The discrepancy in the immunostaining between the two studies could be due to staining or imaging conditions, or due to the points in migration assessed. Similar to the previous study (Penfield and Montell, 2023), we find that Lamin B is prominent in the nucleoskeletons of both the border and polar cells. However, by examining multiple points in migration we find the nucleoskeleton changes during migration. Emerin, which was not assess in the previous study (Penfield and Montell, 2023), like Lamin B, in early migration is present within the nucleoskeletons of both the border and polar cells. As migration proceed, Emerin levels progressively decrease in the nucleoskeleton of all the cells within the cluster but higher levels are retained in the polar cells, whereas Lamin B levels progressively increase within the border cells. These findings are consistent with further nuclear softening of the border cells during migration.

We speculate that these changes in the nucleoskeleton of the border cell cluster are, in part, driven by the environment. Supporting this idea, there are two modes of border cell migration (Bianco *et al*., 2007). The initial phase is a polarized and partially mesenchymal-like migration, whereas the late phase is a rotational migration. This change in migration occurs as the border cell cluster reaches the anterior edge of the layer of nurse cells adjacent to the oocyte (Bianco *et al*., 2007). Further, recent work reveals the topography of the nurse cells and the amount of extracellular space has a stereotypically pattern (George *et al*., 2025); this is the environment that the border cells are migrating through and thereby, the environment being sensed and responded to by the border cells. The study found there are two regions where there is more space, where migration slows, before entering constrictions (George *et al*., 2025). Notably, the latter of these spaces coincides with the change in migration mode (Bianco *et al*., 2007). Further, we used the two points where there is more space, or crevasses, to define early, middle and late migration, supporting that the changes in the nucleoskeleton observed likely reflect changes in nuclear response to the environment.

In early and mid-migration, the nucleoskeleton of the border cells exhibits front/back polarity within the cluster. Specifically, Lamin B is high in the nucleoskeleton of the border cells at the front of the cluster, while Emerin is high at the back of the cluster. We speculate that the front/back polarity of the nucleoskeleton allows the front of the cluster to undergo more nuclear deformation (less Emerin), expanding the space between the nurse cells. This increased space allows the trailing border cells to retain stiffer nuclei. Indeed, a recent study supports that as the border cells migrate through the nurse cells, they change the topography and spacing between the nurse cells (George *et al*., 2025). Further, the leading border cells deform more than the trailing cells (Penfield and Montell, 2023). In conjunction with this information, our findings lead to the model that nucleoskeletal changes allow for nuclear softening and deformation of the front of the cluster as migration proceeds and these changes likely facilitate border cell migration.

Multiple lines of evidence support that the nucleoskeleton plays a critical role in border cell migration. Penfield and Montell found, by assessing whether the border cells complete their migration by S10A, that knockdown of Lamin B but not Lamin A in the border cells impaired migration, as did increasing Lamin B levels (Penfield and Montell, 2023). These findings are consistent with our data that Lamin B, but not Lamin A is present in the border cells, and Lamin B is dynamic during border cell migration. They also found that overexpression of Lamin A in the border cells didn’t result in a failure to complete migration by S10A (Penfield and Montell, 2023). Similarly, we find that border cell overexpression of Lamin A does not alter migration during S9. However, this overexpression of Lamin A may not sufficiently upregulate levels within the nucleoskeleton to impair migration due to cellular mechanisms that control Lamin A levels and/or activity, or due to the construct used. Supporting the latter idea, Lamin A overexpression was driven by the UASp promoter, which is optimized for germline expression and is expressed at lower levels in somatic cells (Rorth, 1998; DeLuca and Spradling, 2018). To overcome this limitation, we overexpressed a Lamin A mutant, R237P; this is a UASt construct and is, therefore, able to be highly expressed in somatic cells (Shaw *et al*., 2022). This mutation mimics the human Emery Dreifus Muscular Dystrophy mutation H222P (Bonne *et al*., 2000). Multiple lines of evidence support that H222P (human cell and mouse models) and R327P (Drosophila) retain significant Lamin A function and may even increase function. Specifically, H222P expression in micro-constricted muscle cells increases cellular stiffness, likely to due to increased nuclear stiffness, compared to cells overexpressing wild-type Lamin A (Chatzifrangkeskou *et al*., 2020). Further, in mouse models, H222P expression in primary myoblasts did not alter differentiation, exhibit the age-dependent contractility decline, or increase cell death – phenotypes observed with other disease mutations or Lamin A knockout (Earle *et al*., 2020). Further, in Drosophila larval muscle, R237P overexpression in the context of wild-type Lamin A results in overly rounded nuclei, suggesting its presence increases nuclear stiffness (Shaw *et al*., 2022). Further, when R237P is expressed Lamin A localizes primarily to the nucleoskeleton (both mutant and wild-type), expression of R237P did not impact Lamin B localization or levels, and the nucleoskeleton and cytoskeleton remained mechanically coupled (Shaw *et al*., 2022). We find that overexpression of Lamin A R237P results in delayed border cell migration during S9. This finding is consistent with the model that nucleoskeletal remodeling, and likely nuclear softening, facilitates border cell migration.

### PGs regulate nucleoskeletal changes during collective migration

Both nucleoskeletal remodeling and polarity during border cell migration are dependent on PG signaling. When PG signaling is lost, Lamin A is prevalent within the nucleoskeleton of both the border and polar cells, Emerin remains high in the nucleoskeleton of the border cells, and Lamin B decreases in nucleoskeleton of the border cells throughout migration. These alterations in nucleoskeletal composition suggests the border cell nuclei are stiffer and less deformable in the absence of PG signaling. Further, Lamin B and Emerin levels lack striking front/back polarity within the cluster when PG signaling is lost, suggesting that there is no longer a softening of the nuclei at the front of the cluster. We hypothesize these changes contribute to the delayed border cell migration during S9 in *dCOX1* mutants. Together these data lead to the model that PG signaling promotes nucleoskeletal remodeling and polarity necessary for the on-time, collective migration of the border cells.

It is likely this mechanism is used in other cell-types and organisms, as PG signaling promotes invasive cell migration across contexts. In zebrafish, COX activity and PGE_2_ signaling are required for cell migration during gastrulation (Cha *et al*., 2005; Cha *et al*., 2006). PG signaling is also co-opted in cancer to drive increased migration and invasion (Singh *et al*., 2005; Kim *et al*., 2010; Fujino *et al*., 2011; Li *et al*., 2012; Vo *et al*., 2013; Wu *et al*., 2017; Demirkol Canli *et al*., 2020). While there is extensive evidence that PG signaling promotes invasive cell migration, there is almost no data connecting PG signaling to the nucleoskeleton in any context. The strongest connection comes from a study on chondrocytes, where PGE_2_ signaling drives Lamin A accumulation; this is not in a cell migration context, but contributes to osteoarthritis (Attur *et al*., 2012). Further, in leukemia, PGE_2_ signaling regulates Lamin B cleavage to regulate cell death (Shehzad *et al*., 2015). In muscle differentiation, Notch signaling prevents differentiation by activating PGE_2_ signaling, and while not directly connected to PG signaling, differentiation requires increased Lamin A (Sakai-Takemura *et al*., 2020). Therefore, this study is the first to connect PG signaling to regulating the nucleoskeleton during cell migration.

How does PG signaling regulate nucleoskeletal dynamics to facilitate border cell migration? One possibility is that PG signaling is required to facilitate mechanotransduction to the nucleus. Such mechanotransduction begins with cell adhesions and cell surface receptors sensing environmental forces and transmitting those forces to the actin cytoskeleton. One key mechanosensitive cell adhesion receptor class is integrins (Huttenlocher and Horwitz, 2011). While the precise functions of integrins in border cell migration are unclear (Dinkins *et al*., 2008; Llense and Martin-Blanco, 2008), we find that PG signaling is required for integrin enrichment on the border cells, as loss results in dispersed integrin localization (Fox *et al*., 2020; Mellentine *et al*., 2023). Such a change could reduce mechanotransduction from outside the cell to the cytoskeleton. The cytoskeleton then needs to transmit the mechanical information to the nucleus. A common way to achieve this is by the LINC Complex, which attaches to the actin and microtubule cytoskeletons, transverses the nuclear envelope and interacts with the nuclear lamina (Mejat and Misteli, 2010; Chang *et al*., 2015). Whether the LINC Complex has a role in border cell migration remains unclear. RNAi knockdown of LINC Complex components in the border cells did not result in delamination defects or a failure to complete border cell migration by S10A (Penfield and Montell, 2023). The impact of knockdown on migration during S9 has not been reported and the genes encoding the LINC Complex have multiple splice forms that are not fully targeted by the RNAi lines; thus, it remains possible that the LINC Complex contributes to border cell migration. One adaptor connecting the LINC Complex to the actin cytoskeleton is Fascin. In this context Fascin, which is best known as an actin bundling protein, is required for invasive single cell migration of mammalian cells *in vitro* (Jayo *et al*., 2016). We find that Fascin localizes perinuclearly in the Drosophila nurse cells, and this localization is both LINC Complex-and PG signaling-dependent, supporting this role of Fascin is conserved in flies (Groen *et al*., 2015; Jayo *et al*., 2016). Notably, Fascin is required for on-time border cell migration (Lamb *et al*., 2020; Lamb *et al*., 2021) and is a downstream effector of PG signaling during border cell migration (Fox *et al*., 2020). Thus, PG signaling may promote Fascin’s LINC Complex role to facilitate force transmission to the nucleus to drive the nucleoskeletal changes required for on-time border cell migration.

There are multiple alternative possibilities for how PGs could regulate the nucleoskeleton within the border cell cluster. For example, PG signaling may induce nucleoskeletal changes within the border cell cluster by altering the environment. Changes in environmental stiffness can be relayed, as described above, to the nucleus. Loss of PG signaling results in both increased border cell cluster and nurse cell stiffness, as observed by increased active non-muscle myosin II (subsequently referred to as myosin) at the cell cortices (Mellentine *et al*., 2023). Notably, knockdown of Fascin, a downstream effector of PGs, in the border cells results in similar increases in stiffness (Lamb *et al*., 2021). In other systems, Fascin’s actin bundling activity limits myosin activity (Elkhatib *et al*., 2014). Therefore, PG signaling may promote Fascin-dependent actin bundling to control both border cell and nurse cell/environmental stiffness, which is sensed by the border and polar cell nuclei to drive nucleoskeletal changes necessary for border cell migration. Another mechanism whereby PGs may regulate the nucleoskeleton is by directly signaling to alter the post-translational modifications of the lamins (Sharma *et al*., 2025); such modifications could lead to Lamin A network disassembly and a loss of Emerin on the inner nuclear envelope. It is likely that PG regulation of the nucleoskeleton is complex and involves more than one of these (and likely other) mechanisms. Future studies are needed to uncover how PGs regulate nucleoskeletal dynamics during cell migration.

Intriguingly, mechanotransduction to the nucleus may also control the production of PGs. Cellular confinement leads to the unfolding and stretching of the nucleus. These changes activate nuclear localized cytosolic phospholipase A2 to produce arachidonic acid (Enyedi *et al*., 2016; Lomakin *et al*., 2020; Venturini *et al*., 2020). Arachidonic acid is the substrate for PG production (Funk, 2001; Tootle, 2013). In zebrafish and HeLa cells this confinement-dependent release of arachidonic acid regulates myosin activity (Enyedi *et al*., 2016; Lomakin *et al*., 2020; Venturini *et al*., 2020). As discussed above, PG signaling limits myosin activity in both the border cells and the nurse cells (Mellentine *et al*., 2023). Thus, there may be a positive feed forward pathway whereby PG signaling regulates nucleoskeletal dynamics and nuclear deformation, and this, in turn, promotes additional PG synthesis and signaling; together, this feed-forward pathway may promote on-time border cell migration.

## Conclusion

In summary, this study provides the first *in vivo* evidence that the nucleoskeleton exhibits dynamic changes and polarity that facilitate a collective migration. It also identifies a novel means of regulating the nucleoskeleton during collective migration – PG signaling. Given the conservation of nucleoskeletal components, mechanotransduction machinery, and PG signaling, these same mechanisms are likely used to facilitate cell migration across organisms and contexts.

## Materials and Methods

### Reagents and Resources

See Supplementary Table S1 for the specific genotypes used in each figure/panel and Supplementary Table S2 for detailed information on reagents used in these studies. Supplementary Table S3 includes all raw data used in this study.

### Fly Stocks

Fly stocks were maintained on Bloomington standard fly food at room temperature, except where noted. Stocks used were *y^1^w^1^*(Bloomington Drosophila Stock Center (BDSC), #1495), *P{GAL4-slbo.2.6}3/TM6B, Tb1* (BDSC, #58435); *UASp Lamin C* (BDSC, #605310); and p*UASt Lamin C R237P (11F-1)* (kind gift from Lori Wallrath; (Shaw *et al*., 2022)). WellGenetics INC (Taipei, Taiwan) generated a new null allele of *dCOX1/Pxt*, *dCOX1^CR2^* (which we refer to as *dCOX1-/-*), using CRISPR/Cas9 mediated repair to excise the complete *dCOX1/Pxt* gene in the *w^1118^* recipient strain. Briefly, the cut sites were located +108 nucleotides from the ATG of *dCOX1* (CRISPR Target Site [PAM]: GGTGCAGGAGTCTGCAAATC[AGG]) and -20 nucleotides from the stop codon of *dCOX1*(CRISPR Target Site [PAM]: CAACCTGCCTTCTGTTAACT[TGG]). A cassette encoding a floxed 3xP3-RFP and two homology arms replaced the deleted *dCOX1* gene. Successful integration was confirmed by sequencing. Afterwards the 3xP3-RFP selection marker was removed by Cre-mediated recombination. For overexpression experiments, crosses were kept at room temperature and progeny were kept at 25°C for 3 days prior to immunofluorescence experiments.

### Immunofluorescence

Before immunofluorescence staining, newly eclosed flies were fed wet yeast paste every day for 2-4 days. Drosophila ovaries (10-15 pairs per sample) were dissected into room temperature Grace’s insect medium (Corning, Corning, NY). Ovaries were fixed for 10 min using 4% paraformaldehyde diluted in Grace’s medium. Samples were washed six times for 10 min each at room temperature in antibody wash (1X phosphate-buffered saline [PBS], 0.1% Triton X and 0.1% bovine serum albumin [Sigma-Aldrich, Burlington, MA]) Primary antibodies were diluted in antibody wash and incubated overnight at 4°C with rocking. The following monoclonal antibodies were obtained from the Developmental Studies Hybridoma Bank (DSHB), created by the NICHD of the NIH and maintained at The University of Iowa, Department of Biology, Iowa City, IA: mouse anti-Lamin C 1:250 (LC28.26, RRID: AB_528339; (Riemer *et al*., 1995)) and mouse anti-Lamin Dm0 1:200 ( ADL101, RRID: AB_528332; (Riemer *et al*., 1995)). Rabbit anti-dEmerin/Otefin was used at 1:1000. This antibody was generated by GenScript against a peptide of the first 187 amino acids of dEmerin/Otefin, similar to what was previously described (Barton *et al*., 2014). After incubation in primary antibody, samples were washed six times, 10 min each in antibody wash. Samples were then incubated in secondary antibodies overnight at 4°C with rocking. The following secondaries were used at 1:500: AF488T goat anti-mouse (RRID: AB_2534069; Thermo Fisher Scientific, Waltham, MA), AF568T goat anti-rabbit (RRID:AB_10563566; Thermo Fisher Scientific). Alexa Fluor 647-conjugated phalloidin (A12380 and A22287; Thermo Fisher (Invitrogen); Waltham, MA) diluted 1:250 was included in both primary and secondary antibody incubations. After secondary antibody incubation, the samples were washed six times, 10 min each in antibody wash. 4′,6-diamidino-2-phenylidole (DAPI; 5 mg/mL; D9542; Millipore Sigma; Burlington, MA) staining was performed at a concentration of 1:5000 in 1X PBS for 10 min. Samples were then rinsed in 1X PBS and mounted on slides in 1 mg/ mL phenylenediamine in 50% glycerol, pH 9 (Platt and Michael, 1983) or Vectashield (RRID:AB_2336789; Vector Laboratories, Newark, CA). A minimum of three independent experiments were performed for each staining.

### Image acquisition and processing

Microscope images of stained and fixed Drosophila follicles were obtained using a Leica DM8 Stellaris confocal microscope (Leica Microsystems) using either a HC PL APO CS 20x 0.75 dry objective at a 1.40 zoom or a HC PL APO CS2 63x/1.40 oil objective at 3.50 zoom. To ensure fluorescent intensities could be quantified and compared, for a given experiment, the microscope settings were identical for each genotype in said experiment and set to ensure there were not overexposed pixels based on the genotype with the highest stain intensity. Images were rotated and cropped, and scale bars were added using FIJI (RRID: SCR_002285; (Abramoff *et al*., 2004)). As indicated in the figure legends, Adobe Photoshop was used to brighten images. Adobe Illustrator was used to assemble figures (Adobe, RRID: SCR 010279; San Jose, CA).

### Image quantifications

Each S9 follicle was defined as being at early, middle, or late migration. The two positions within the follicle where eight nurse cells come together to create a crevasse (George *et al*., 2025) were used to define early, middle, and late migration. Delamination at the anterior end of the follicle to immediately preceding the first nurse cell crevasse was defined as early migration. The first crevasse to immediately preceding the second crevasse was defined as middle migration. The second crevasse to when the border cells reach the oocyte was defined as late migration.

To quantify fluorescence intensity, Imaris software (RRID:SCR_007370; Zurich, Switzerland) was used to create a 3D rendering of 63x confocal image stacks. A surface was created around the border cell cluster and cut so that each individual nuclei was contained in a separate surface. The nuclei were designated as either polar or border cells. The fluorescent mean intensity was measured for each nuclei. We averaged the fluorescent mean intensity of the border cells and the polar cells for an individual border cell cluster. Then the relative fluorescence intensity (RFI) was measured by dividing the average fluorescent mean intensity of the border cells by the averaged fluorescent mean intensity of the polar cells for each cluster. To compare fluorescent intensity of polar cells and border cells within the same cluster we averaged the fluorescent mean intensity of the two polar cells for each cluster and averaged the fluorescent mean intensity of the border cells for each cluster. To compare fluorescent intensity between *wild-type* and *dCOX1-/-*, we averaged the fluorescent mean intensity of the two polar cells and the border cells for each cluster.

To assess the nucleoskeletal polarity within the border cell cluster, confocal image stacks were processed as described above and the mean fluorescent intensity was measured for each border cell. The polar cells were used as a reference to determine which border cells were designated as front or back cells. Border cells were designated as front cells if they were predominantly posterior to the polar cells (closest to the oocyte) and as back cells if they were predominantly anterior to the polar cells. In some cases, the border cells were in line with the polar cells; we designated these border cells as side cells and omitted them from the quantification. The fluorescent mean intensity was measured for each nuclei. The fluorescent mean intensities of the front border cells and the back border cells for each individual border cell cluster were averaged. The average mean fluorescent intensity of the back border cells was subtracted from the average mean fluorescent intensity of the front border cells for each individual border cell cluster to generate a polarity score. A negative value indicates that the nucleoskeletal protein is more prominent in the front of the cluster, whereas a positive value indicates the nucleoskeletal protein is more prominent at the back of the cluster.

Data were compiled using Microsoft Excel (Microsoft, RRID: SCR_016137 Redmond, WA). Graphs were generated and statistical analyses performed using Prism 9 (GraphPad Software RRID: SCR_ 002798). Robust nonlinear regression and outlier tests were performed on all quantifications and outliers were omitted from the assessments. A minimum of three independent experiments were performed for each stain and genotype.

### Quantification of border cell migration, cluster length, and border cell number

Quantification of the Migration Index was performed as described previously (Fox et al., 2020; Lamb et al., 2020). Briefly, measurements of S9 follicles were performed using Fiji software (Abramoff *et al*., 2004) on maximum projections of 2-4 confocal slices of follicles stained for Lamin A, Lamin B, or Emerin. A line segment was used to measure the distance in microns from the anterior end of the follicle to the leading edge of the border cell cluster; this was defined as the border cell distance. Another line segment was used to measure the distance from the anterior end of the follicle to the anterior end of the outer follicle cells: this was defined as the outer follicle cell distance. The entire follicle length was also measured along the anterior-posterior axis. The migration index was calculated by dividing the border cell distance by the follicle cell distance. Cluster length was determined by measuring the distance from the rear to the front of the border cell cluster (nondetached border cell clusters were not included in the analysis). The number of border cells per cluster was quantified during Imaris analyses; depending on the experiment the border cells were identified with either Lamin A, Lamin B, or Emerin staining. Data were compiled in and calculations were performed in Excel, and graphs were generated and statistical analyses performed using Prism 9. A minimum of three independent experiments were performed for quantifications.

### Western Blots

Whole ovary pairs (5 total per sample) were dissected at room temperature in Grace’s medium and transferred to a 1.5 ml tube containing 50 µl 1xPBS. 50 µl of 2x Laemmli buffer was added and lysis was performed by grinding tissue with plastic pestles (RNase-free disposable pellet pestles; Thermo Fisher Scientific). Samples were boiled and briefly spun down and stored at - 20°C until use. Samples were again boiled for 10 min and briefly spun down before quantifying protein and loading onto gel. Protein amount was quantified using a Pierce 660-nm Protein Assay (Thermo Fisher Scientific). Briefly this assay works by mixing Pierce 660-nm Protein assay reagent (Thermo Fisher Scientific) and 25mg/mL of Neutralizer (G-Biosciences, St. Louis, MO) with BSA protein standards (0µg/mL-2500µg/mL) and protein samples. Standards and samples were analyzed on a Synergy LX plate reader (Agilent Technologies, Santa Clara, CA). Prism 9 was used to generate a standard curve and extrapolate protein sample amounts. 20 µg/mL of protein was loaded per sample. Western blots were performed using standard methods. Briefly, samples were run on 10% SDS-PAGE gels and transferred onto nitrocellulose membranes (Amersham Protran 0.2 µm NC; GE Healthcare Life Sciences, Marlborough, MA). The ladder used was Precision Plus Protein Dual Color Standards (161-0374; Bio-Rad Laboratories, Hercules, CA). Blot was cut horizontally at ∼75 kDa prior to primary antibody incubation to allow for the assessment of two different proteins. Blots were incubated for 30 min in Western blot block (5% non-fat dry milk in 1x Tris-Buffered Saline with 0.1% Tween 20 [TBST]). The following primary antibodies and concentrations were used: rabbit anti-dCOX1 1:5000 (Spracklen *et al*., 2014) and mouse anti-alpha tubulin 1:500 (DSHB, RRID: AB_1157911; (Jerka-Dziadosz *et al*., 1995)). Antibodies were diluted in Western blot block and incubated overnight at 4°C. After primary antibody incubation, blots were washed three times in 1x Tris-Buffered Saline (TBS) and once in 1x TBST for 10 min each. Blots were then incubated in secondary antibody for 2.5 hours at room temperature. Following secondary incubation, blots were washed three times with 1x TBS and twice with 1x TBST, for 10 min each. The following secondaries were used at 1:5000 Western blot block: Peroxidase-AffiniPure Goat Anti Rabbit IgG (H+L) and Peroxidase-AffiniPure Goat Anti Mouse IgG (H+L) (Jackson ImmunoResearch Laboratories, West Grove, PA). Blots were developed with SuperSignale West Pico Chemiluminescent Substrate (Thermo Scientific) and imaged using the Li-Cor Odyssey FC (Li-Cor, Lincoln, NE). Three independent experiments were performed.

## Supporting information

STable 1

STable 2

STable 3

## Acknowledgements

We thank Julia Dorale for data processing, and the Tootle lab for helpful discussions and careful review of the manuscript. Stocks obtained from the Bloomington Drosophila Stock Center (NIH P40OD018537) were used in this study. FlyBase (release FB2024_04, NIH U41 HG000739) was used for information on stocks. At the University of Iowa, Information Technology Services – Research Services provided data storage support.

## Data and resource availability

All relevant data and details of resources can be found within the article and its supplementary information.

## Abbreviations

PG: prostaglandin
S8: Stage 8
S9: Stage 9
LINC: Linker of the Nucleoskeleton and Cytoskeleton

