## Supplemental Figures for "Prostaglandins regulate the nucleoskeleton during Drosophila border cell migration"

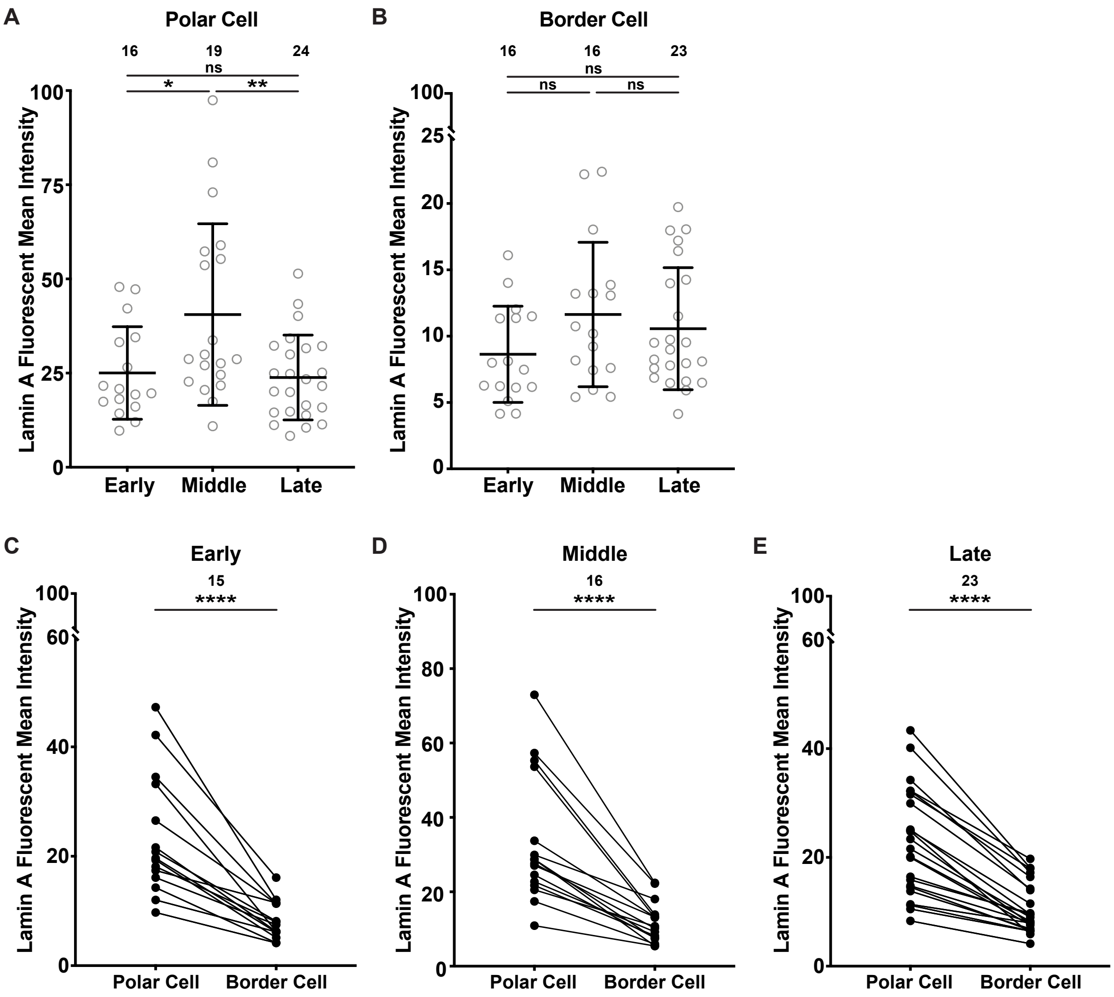


**Supplementary Figure 1: Lamin A is predominantly within the nucleoskeleton of the polar cells and exhibits little change during border cell migration. (A-B)** Graphs of Lamin A mean fluorescent intensity in the nucleoskeletons of the polar cells (**A**) or border cells (**B**) at the indicated points in migration. Circle = individual BC cluster; *n* = number of follicles, lines = averages and error bars = standard deviation (SD). ns > 0.05, *p<0.05, ** p < 0.01, unpaired *t*-test, two-tailed. **(C-E)** Graphs of paired mean fluorescent intensity of Lamin A in the nucleoskeletons of the polar cells and border cells within each cluster at early (**C**), middle (**D**) and late (**E**) migration. Circle = individual BC cluster; *n* = number of follicles, lines = polar cells and border cells from the same cluster. **** p < 0.0001, paired *t*-test, two-tailed. The level of Lamin A within the nucleoskeletons of the polar cells and border cells remains largely constant throughout migration, with the exception that there is a slight decrease in the polar cells during late migration (**A-B**). However, Lamin A is significantly higher in the nucleoskeleton of the polar cells compared to the border cells within each cluster throughout migration **(C-E)**.

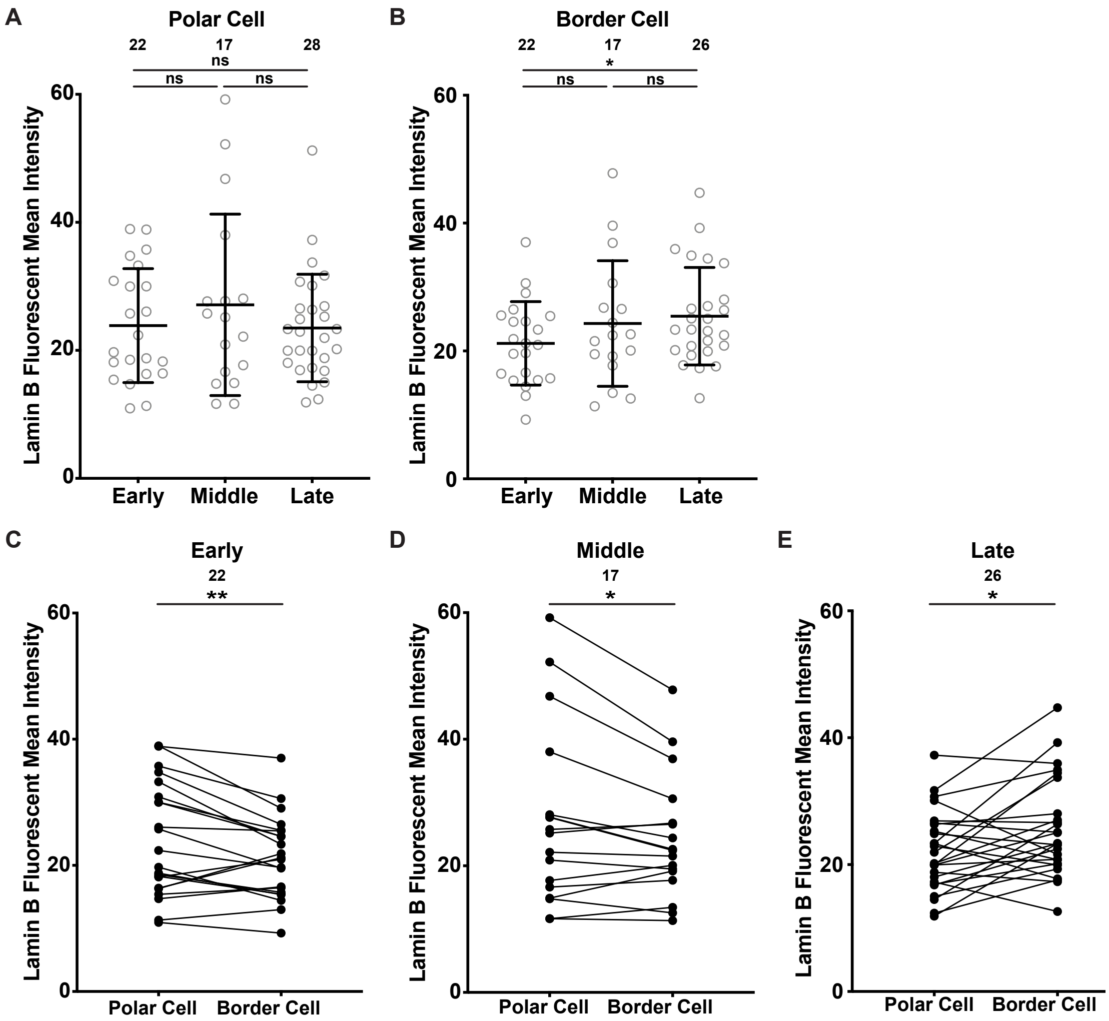


**Supplementary Figure 2: Lamin B increases in the nucleoskeleton of the border cells during border cell migration. (A-B)** Graphs of Lamin B mean fluorescent intensity in the nucleoskeletons of the polar cells (**A**) or border cells (**B**) at the indicated points in migration. Circle = individual BC cluster; *n* = number of follicles, lines = averages and error bars = standard deviation (SD). ns > 0.05, * p < 0.05, unpaired *t*-test, two-tailed. **(C-E)** Graphs of paired mean fluorescent intensity of Lamin B in the nucleoskeletons of the polar cells and border cells within each cluster at early (**C**), middle (**D**) and late (**E**) migration. Circle = individual BC cluster; *n* = number of follicles, lines = polar cells and border cells from the same cluster. *p<0.05, ** p < 0.01, paired *t*-test, two-tailed. Lamin B is present at a consistent level within nucleoskeleton of the polar cells throughout migration (**A**), whereas Lamin B levels within the border cells increase in late migration (**B**). Within individual border cell clusters, Lamin B is generally higher in the nucleoskeleton of the polar cells compared to the border cells in early and mid-migration but is higher in the border cells in late migration (**D-E**).

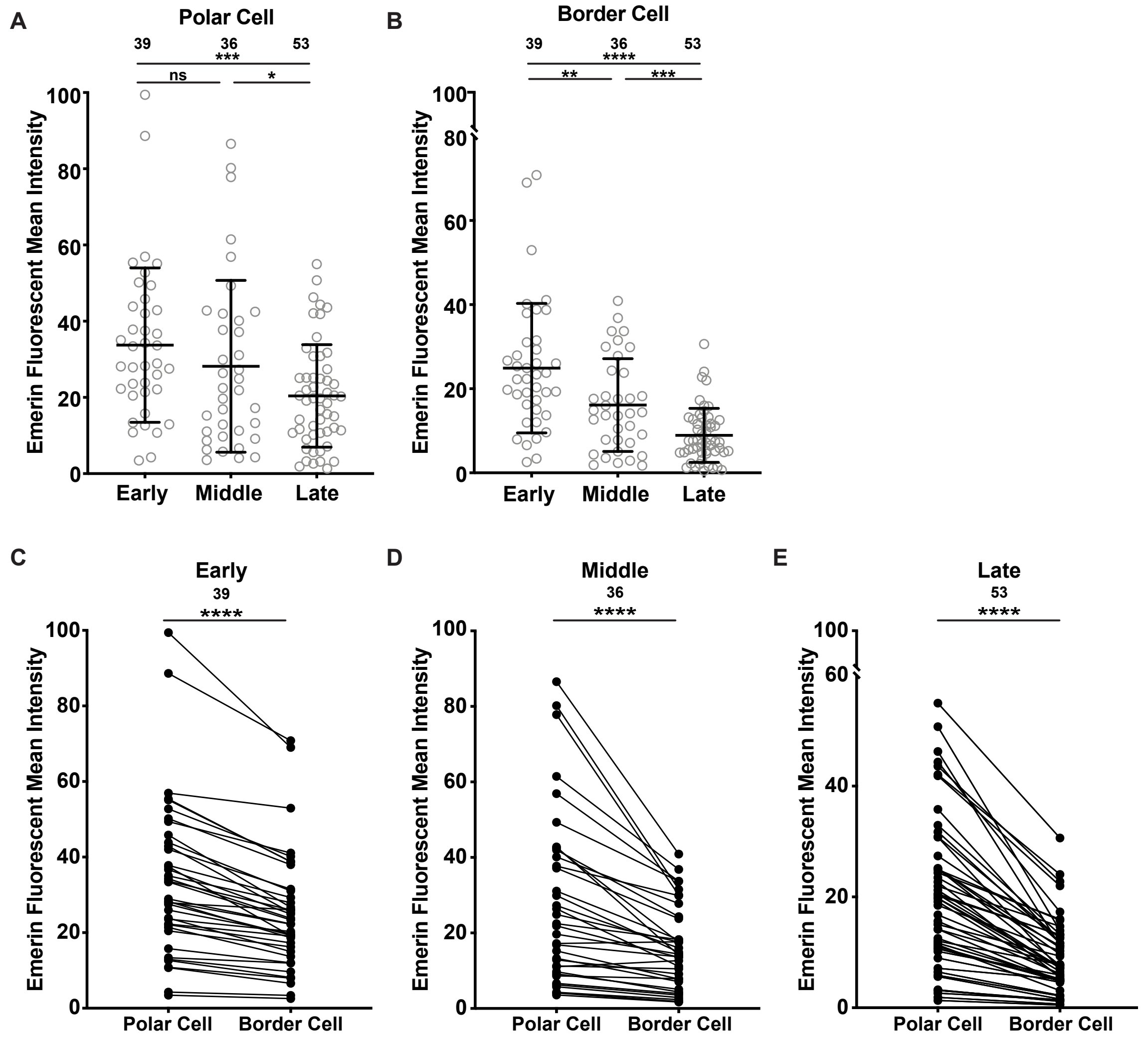


**Supplementary Figure 3: Emerin decreases in the nucleoskeletons of the polar cells and border cells during migration. (A-B)** Graphs of Emerin mean fluorescent intensity in the nucleoskeletons of the polar cells **(A)** or border cells **(B)** at the indicated points in migration. Circle = individual BC cluster; *n* = number of follicles, lines = averages and error bars = standard deviation (SD). ns > 0.05, * p < 0.05, ** p < 0.01, ***p<0.001, unpaired *t*-test, two-tailed. **(C-E)** Graphs of paired mean fluorescent intensity of Emerin in the nucleoskeletons of the polar cells and border cells within each cluster at early (**C**), middle (**D**) and late (**E**) migration. Circle = individual BC cluster; *n* = number of follicles, lines = polar cells and border cells from the same cluster. **** p < 0.0001, paired *t*-test, two-tailed. Emerin levels in the nucleoskeletons of both the polar cells and border cells decrease from early to late migration, this decrease is more striking in the border cells (**A-B**). Within individual border cell clusters, Emerin is more prevalent in the nucleoskeleton of the polar cells compared to the border cells throughout migration (**D-E**).

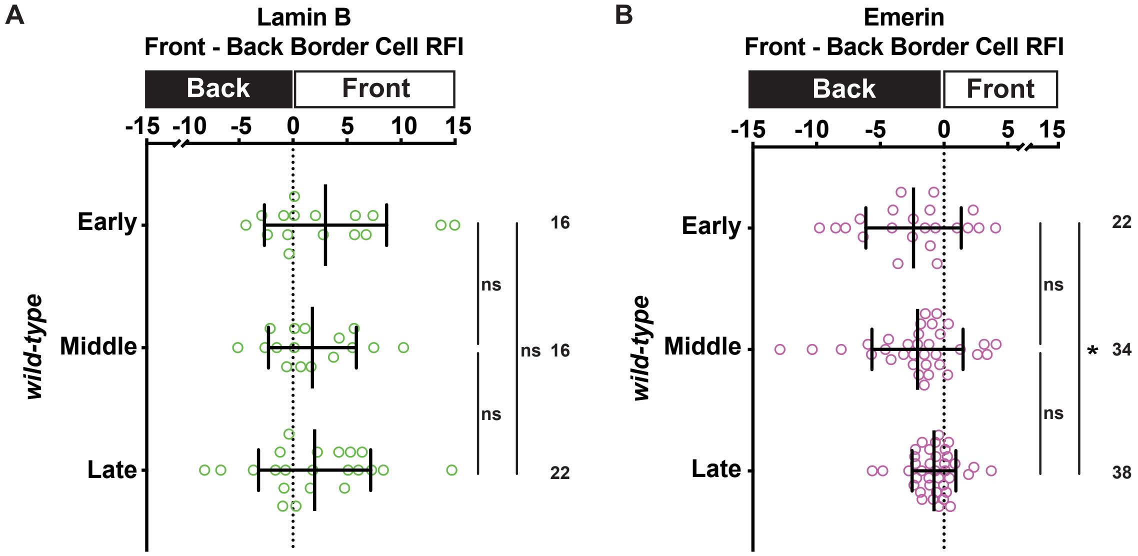


**Supplementary Figure 4: Lamin B and Emerin are polarized within the border cell cluster throughout migration. (A-B)** Graphs of front - back relative fluorescent intensity (RFI) ratio for Lamin B (**A**) and Emerin (**B**) in the nucleoskeleton of the border cells at early, middle, and late migration. Circle = individual BC cluster; *n* = number of follicles, lines = averages and error bars = SD. ns > 0.05, * p < 0.05, unpaired *t*-test, two-tailed. A value of 0 indicates a lack of polarity, indicated by the dotted line. Positive value indicates the nucleoskeletal protein is more prominent at the front of the cluster. Negative value indicates nucleoskeletal protein is more prominent at the back of the cluster. Lamin B is prominent at the front of the cluster throughout migration (**A**). Whereas Emerin is prominent at the back of the cluster throughout migration (**B**); this polarity is less striking during late migration, due to reduction in Emerin in the nucleoskeleton of the border cells during late migration (see SFig. 3).

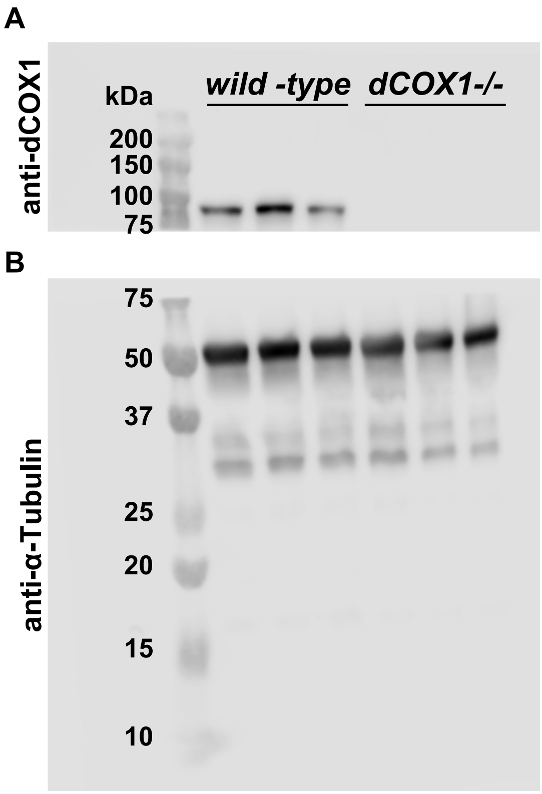


**Supplementary Figure 5: Western blot for dCOX1 expression. (A-B)** Western blot of whole ovary lysates of *wild-type* and *dCOX1-/-*. Blot was cut after transfer and stained with anti-dCOX1 **(A)** or anti-$\alpha$-Tubulin (**B**). Molecular weight ladder in kDa is indicated on the blot. The CRISPR dCOX1 allele is a null **(A-B)**.

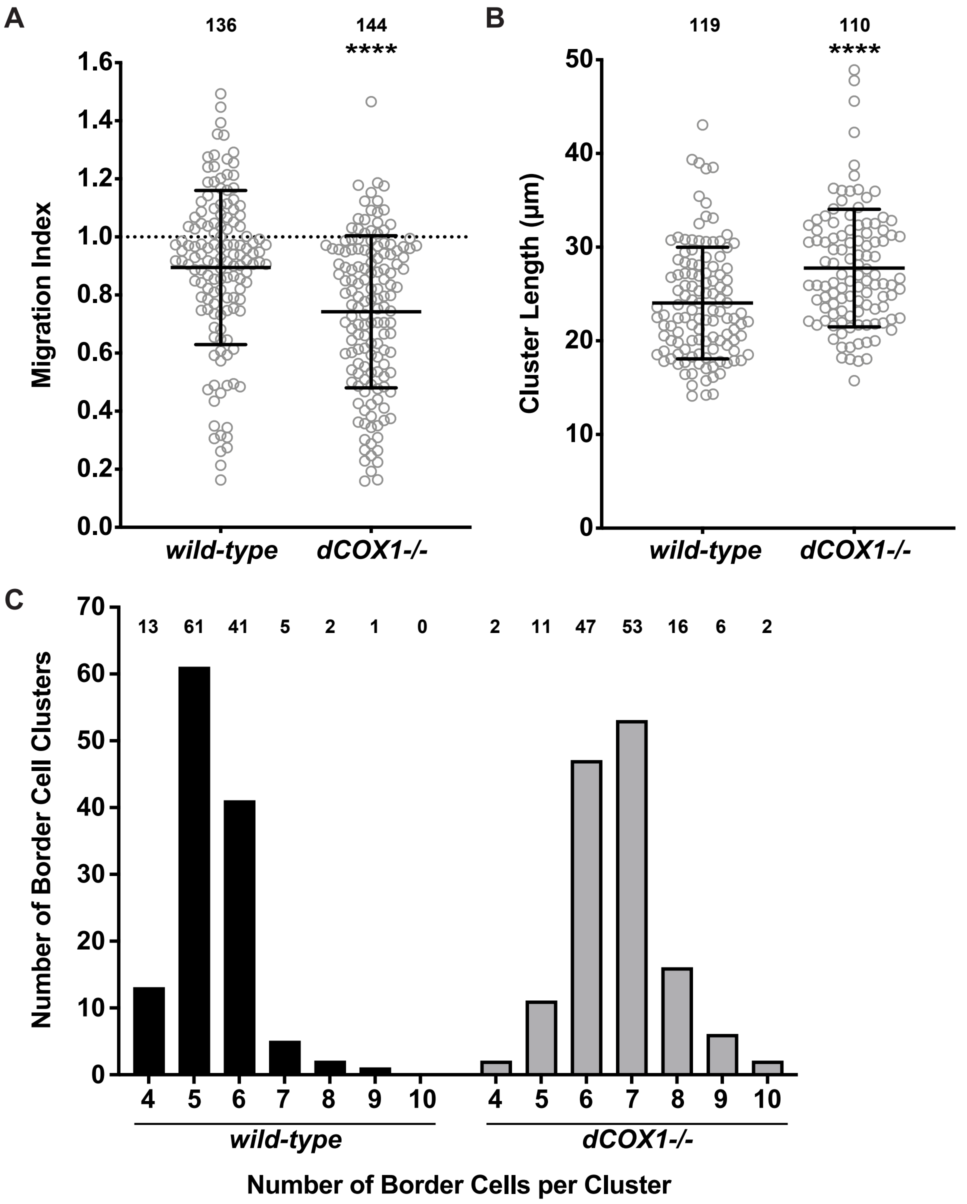


**Supplementary Figure 6: The CRISPR null allele of *dCOX1* recapitulates the border cell defects seen with prior alleles. (A-B)** Graphs quantifying the migration index (**A**) and border cell cluster length (**B**) for wild-type and *dCOX1-/-*. In **A**, on-time border cell migration is indicated by the dotted line at 1. Circle = individual BC cluster; *n* = number of follicles, lines = averages and error bars = SD. **** $p < 0.0001$ unpaired *t*-test, two-tailed. (**C**) Histogram of the number of border cells per cluster for wild-type and *dCOX1-/-*. *n* = number of follicles. ***$p<0.001$, Fisher exact test. Loss of PG signaling results in delayed migration (**A**), elongated clusters (**B**), and an increased number of border cells per cluster (**C**).

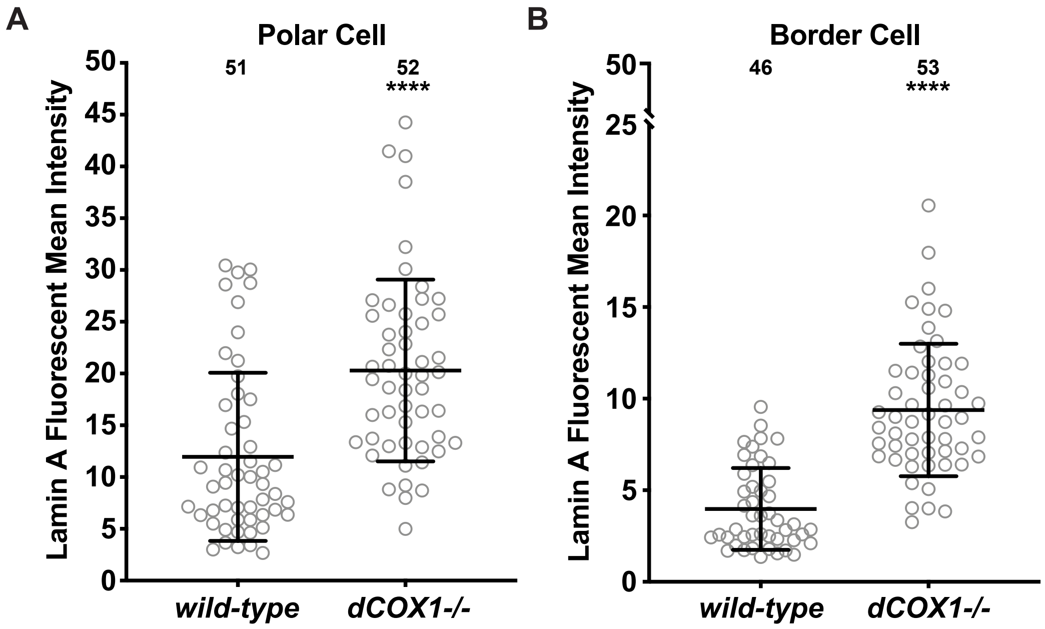


**Supplementary Figure 7: PG signaling normally limits Lamin A levels within the nucleoskeletons of the polar and border cells. (A-B)** Graphs of Lamin A mean fluorescent intensity in the nucleoskeletons of the polar cells (**A**) or border cells (**B**) for wild-type and *dCOX1-/-*. Circle = individual BC cluster; *n* = number of follicles, lines = averages and error bars = standard deviation (SD). **** $p < 0.0001$, unpaired *t*-test, two-tailed. Loss of PG signaling results in an increase of Lamin A in the nucleoskeleton of the polar cells and border cells (**A-B**).

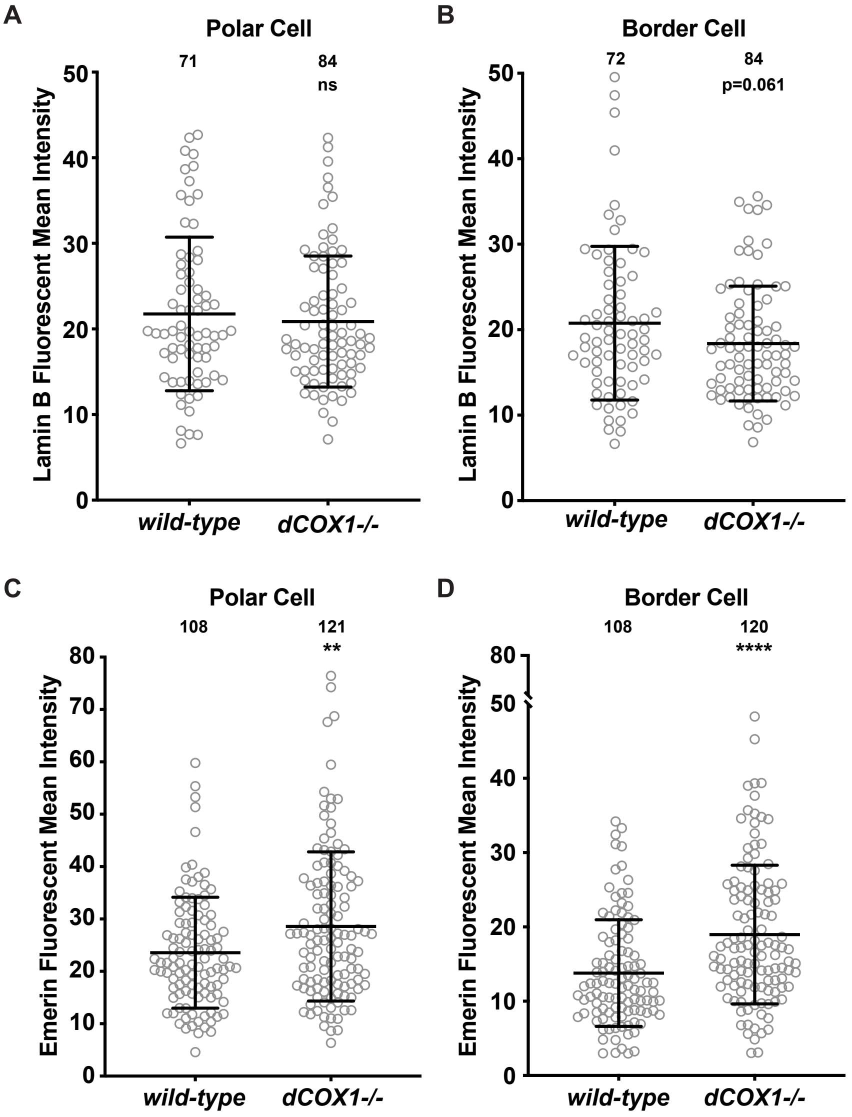


**Supplementary Figure 8: PG signaling limits Emerin levels within the nucleoskeletons of both the polar and border cells, but has a minor role in modulating Lamin B. (A-D)** Graphs of Lamin B (**A-B**) and Emerin (**C-D**) mean fluorescent intensity in the nucleoskeletons of the polar cells **(A, C)** or border cells **(B, D)** for wild-type and *dCOX1-/-*. Circle = individual BC cluster; *n* = number of follicles, lines = averages and error bars = standard deviation (SD). ns > 0.05, ** p < 0.01, **** p < 0.0001, unpaired *t*-test, two-tailed. Loss of PG signaling has no effect on Lamin B levels in the nucleoskeleton of the polar cells (**A**), but results in a slight reduction in the border cells (**B**). Conversely, loss of PG signaling results in increased levels of Emerin in the nucleoskeletons of both the polar and border cells (**C-D**).

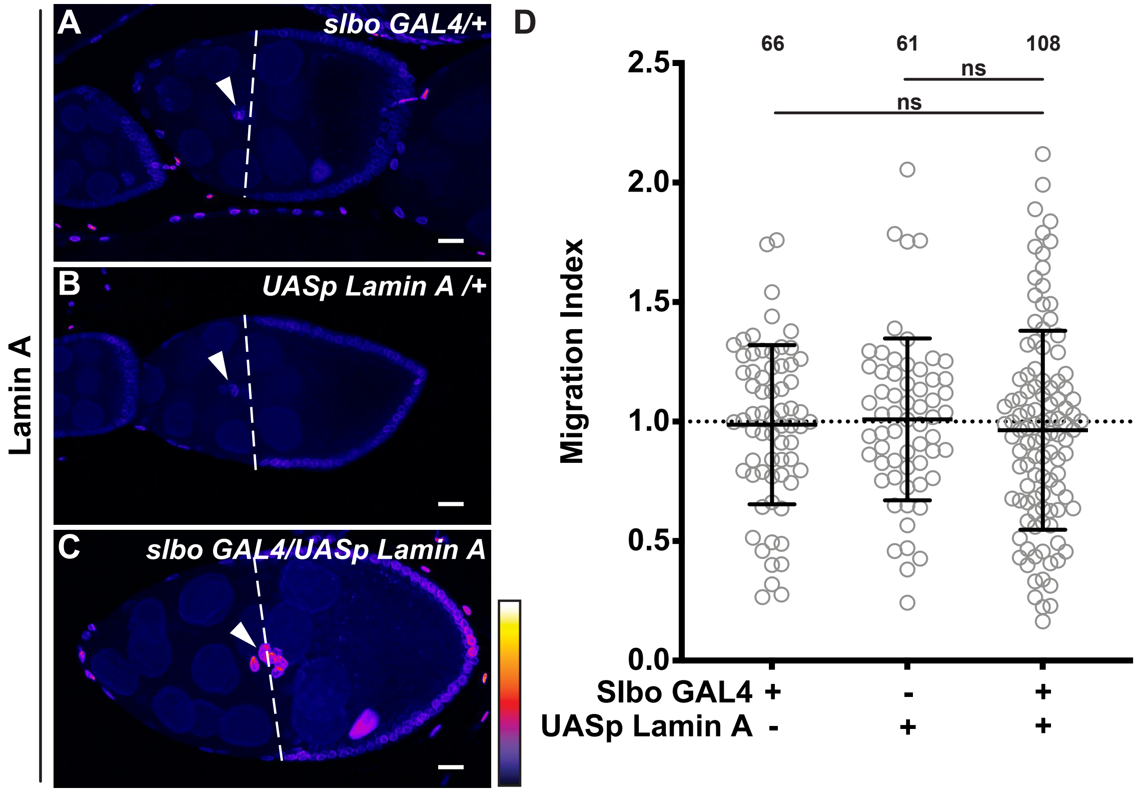


**Supplementary Figure 9: Overexpression of Lamin A does not impact BC migration. (A-C)** Maximum projection of confocal slices of S9 follicles stained for Lamin A in Fire LUT to reflect intensity of the following genotypes: *slbo GAL4/+* (**A**); *UASp Lamin A/+* (**B**); and *slbo GAL4/UASp Lamin A* (**C**). White arrowheads indicate border cell clusters and white dashed lines indicate the position of the outer follicle cells. Scale bars= 20 µm. Images brightened by 30% to increase clarity. **(D)** Graph of migration index for the indicated genotypes. On-time border cell migrated indicated by dotted line. Circle = individual BC cluster; *n* = number of follicles, lines = averages and error bars = SD. ns>0.05, unpaired *t*-test, two-tailed. Both controls and overexpression of Lamin A in the border cells exhibit on-time border cell migration (**A-D**)

**Supplementary Table 1. Genotypes by figure.** List of genotypes shown in each figure panel.

**Supplementary Table 2. Reagents Table.** List of reagents used in the study.

**Supplementary Table 3. Raw data.** Raw data used for all quantifications presented in both the primary and supplemental figures.
